# Transcriptional Reprogramming of Skeletal Muscle Stem Cells by the Niche Environment

**DOI:** 10.1101/2021.05.25.445621

**Authors:** Felicia Lazure, Rick Farouni, Korin Sahinyan, Darren M. Blackburn, Aldo Hernández-Corchado, Gabrielle Perron, Jiannis Ragoussis, Colin Crist, Theodore J. Perkins, Arezu Jahani-Asl, Hamed S. Najafabadi, Vahab D. Soleimani

## Abstract

Adult stem cells are indispensable for tissue regeneration. Tissue-specific stem cells reside in a specialized location called their niche, where they are in constant cross talk with neighboring niche cells and circulatory signals from their environment. Aging has a detrimental effect on the number and the regenerative function of various stem cells. However, whether the loss of stem cell function is a cause or consequence of their aging niche is unclear. Using skeletal muscle stem cells (MuSCs) as a model, we decouple cell-intrinsic from niche-mediated extrinsic effects of aging on their transcriptome. By combining *in vivo* MuSC heterochronic transplantation models and computational methods, we show that on a genome-wide scale, age-related altered genes fall into two distinct categories regarding their response to the niche environment. Genes that are inelastic in their response to the niche exhibit altered chromatin accessibility and are associated with differentially methylated regions (DMRs) between young and aged cells. On the other hand, genes that are restorable by niche exposure exhibit altered transcriptome but show no change in chromatin accessibility or DMRs. Taken together, our data suggest that the niche environment plays a decisive role in controlling the transcriptional activity of MuSCs, and exposure to a young niche can reverse approximately half of all age-associated changes that are not epigenetically encoded. The muscle niche therefore serves as an important therapeutic venue to mitigate the negative consequence of aging on tissue regeneration.

## Introduction

Aging invariably causes a decline in muscle mass, strength, and regenerative capacity, leading to the onset of muscle-wasting diseases such as sarcopenia in the elderly (Berger and Doherty, 2010; Cruz-Jentoft et al., 2019). This is in part due to the decline in numbers and regenerative function of MuSCs (Shefer et al., 2006). The MuSC pool is maintained in a quiescent state under homeostatic conditions and retains the ability to activate in response to injury in order to repair damaged muscle tissue. These processes of activation, differentiation and self-renewal are highly dependent on niche-derived cues (Mourikis et al., 2012; Sampath et al., 2018). Numerous non-myogenic cells such as fibro-adipogenic progenitors (FAPs), macrophages, and endothelial cells, amongst others, have been shown to be important in maintaining MuSC function (Biferali et al., 2019; Joe et al., 2010). However, the cross-talk between MuSCs and neighboring cells is perturbed in aging, leading to a breakdown in MuSC support from the niche (Ancel et al., 2019; Lukjanenko et al., 2016; Lukjanenko et al., 2019). In addition to neighboring cells, age-related changes in the niche also extend to systemic factors originating from blood circulation as well as structural components such as myofibers and extracellular matrix composition (ECM) (Garg and Boppart, 2016; Gilbert et al., 2010; Li et al., 2019; Oh et al., 2016).

As an additional layer of complexity, the niche environment is also highly dynamic and changes during the course of diseases and during normal physiological aging. Given the known changes that occur in the aging niche, there have been efforts to restore MuSC function by altering various components of the niche. Historically, parabiosis experiments have shown that exposure of aged muscle to young circulatory factors can restore the regenerative capacity of aged muscle tissue (Conboy et al., 2005), demonstrating the importance that extrinsic factors can have on MuSC function. While there is evidence for reversibility of the aging muscle phenotype, other studies provide evidence supporting the presence of irreversible cell-intrinsic changes in aged MuSCs. For example, it has been reported that aged MuSCs have a reduced capacity to engraft into healthy adult mice compared to young counterparts (Bernet et al., 2014; Chakkalakal et al., 2012).

Despite the evidence for both cell-intrinsic and niche-mediated changes occurring in aged MuSCs, models to effectively decouple the two have been lacking. Studying MuSCs within the context of their niche and without *ex vivo* expansion has previously been challenging due to the scarcity of cellular material sufficient for genome-wide analyses. In this study, we use allogeneic stem cell transplantation combined with Switching Mechanism at 5’ End of RNA Template (SMART-Seq) technology to directly quantify the effect of the niche on the MuSC gene expression profile. We determine the genome-wide reversibility of age-related altered gene expression by exposure of MuSCs to the young niche environment. Combined with single-cell RNA-seq, ATAC-seq, and Whole Genome Bisulfite (WGBS) sequencing data, we characterize the transcriptomic and epigenetic profile of genes that are reversible and irreversible by the niche to determine signatures that confer plasticity in gene expression in response to the niche.

## Results

### MuSC Subpopulations Display Different Dynamics of Loss Versus Retention in Aging

To determine the effect of aging on the population dynamics of MuSCs and muscle-resident niche cells, we performed single-cell RNA sequencing of young and aged MuSCs, as well as two important non-myogenic niche populations: macrophages (Cui et al., 2019; Wang et al., 2015) and Fibro/Adipogenic progenitor cells (FAPs) (Lukjanenko et al., 2019) (Figures 1A and S1). In addition to an overall reduction in numbers as seen after Fluorescence Activated Cell Sorting (FACS) (Figures 1B-1C and S1), and UMAP clustering (Figure 1D), analysis of scRNA-seq suggests that MuSCs exhibit an age-related shift in their population diversity (Figures 1E-1F). When comparing young and aged FAPs, the primary effect of aging is an overall expansion in cell number in aging (Figure 1G). Additionally, while FAPs appear to be major contributors to the components of the Extracellular Matrix (ECM), FAPs from aged mice exhibit reduced expression of ECM genes such as *Col1a1* and *Col1a2* among others (Figures 1H, 1L and S2A-S2E). Meanwhile, macrophages do not show a significant change in numbers nor a major shift in their population dynamics during aging (Figure 1I). However, a deregulation in the expression of previously identified macrophage marker genes such as *Msrb1* (Lee et al., 2017), *Plcb1* (Di Raimo et al., 2016) and *Gsr* (Parisi et al., 2018) can be observed, possibly indicating a phenotypic switch during aging (Figures 1J-1K and S2F).

**Figure 1.**
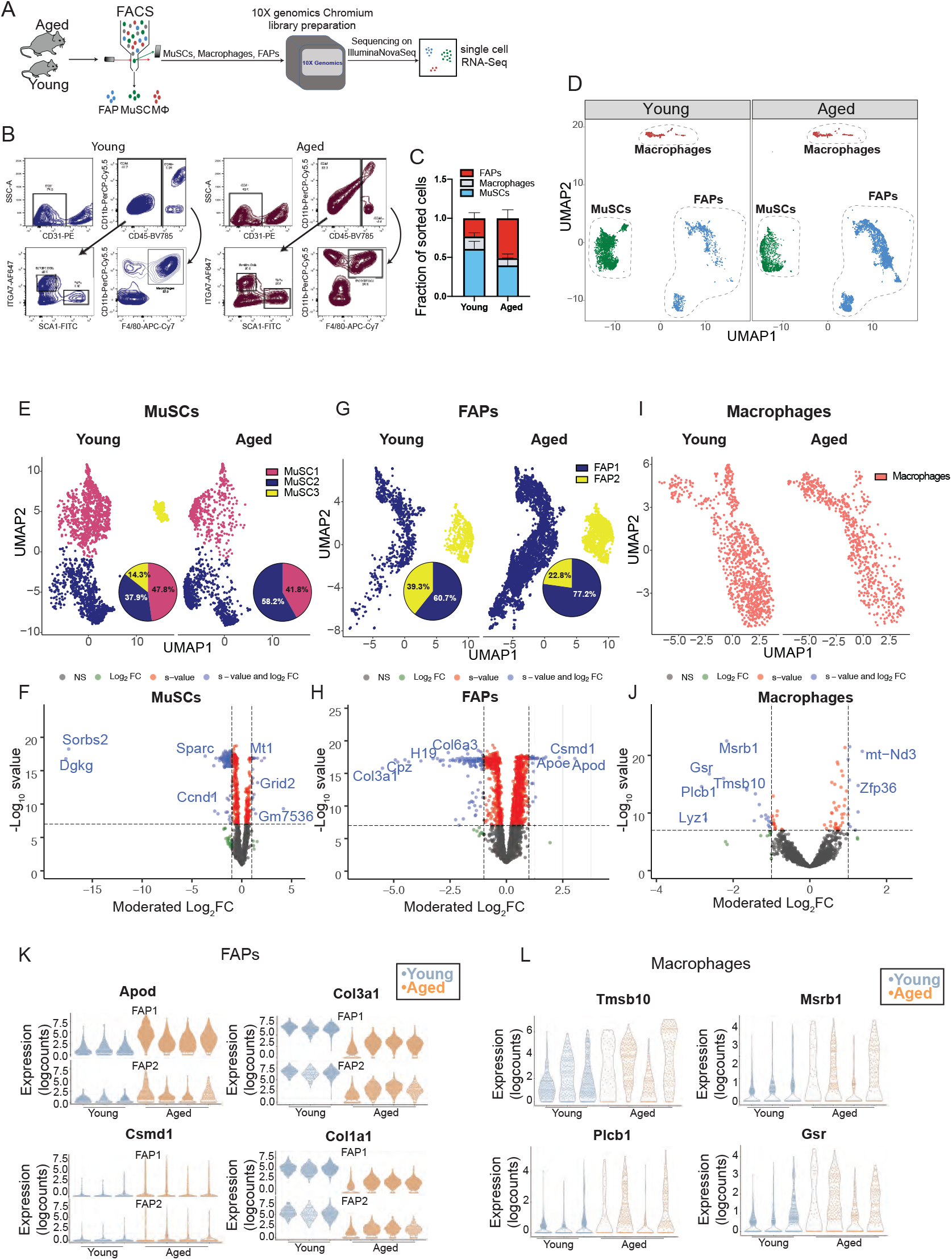
The Population Dynamics of MuSCs and Niche Cells Are Altered in Aging. **(A)** Schematic diagram of the workflow for the isolation and single-cell RNA sequencing of young and aged MuSCs, macrophages and Fibro-Adipogenic Progenitors (FAPs). **(B)** Fluorescence-Activated Cell Sorting (FACS) plots demonstrating the strategy for the simultaneous isolation of MuSCs, macrophages and FAPs from young (5-6 week old) and aged (24 month old) mice. **(C)** Relative proportions of MuSCs, macrophages and FAPs isolated from young and aged mice (n=3 young mice, n=4 aged mice) **(D)** Global UMAP plot of scRNA-seq data from young and aged MuSCs, FAPs, and macrophages (n=3 mice per group) **(E)** UMAP plot of scRNA-seq data from young and aged MuSCs including pie charts showing the proportions of cells in each subpopulation. **(F)** Volcano plots showing genes that are up or down-regulated due to aging in MuSCs at a threshold of moderated LFC >1 and s-value <0.05. **(G)** UMAP plot of scRNA-seq data from young and aged FAPs including pie charts showing the proportions of cells in each subpopulation. **(H)** Volcano plots showing genes that are up or down-regulated due to aging in FAPs at a threshold of moderated LFC >1 and s-value <0.05. **(I)** UMAP plot of scRNA-seq data from young and aged macrophages. **(J)** Volcano plots showing genes that are up or down-regulated due to aging in macrophages at a threshold of moderated LFC >1 and s-value <0.05. **(K)** Violin plots of select representative genes that are differentially expressed between young and aged FAPs. **(L)** Violin plots of select representative genes that are differentially expressed between young and aged macrophages (moderated LFC >1, s-value < 0.05).

Evidence suggests that during aging, cells are lost due to the accumulation of DNA damage and oxidative and replication stress that occur over time (Lombard et al., 2005; Oh et al., 2014). Our data indicates an alternative mechanism is also at play, since different subpopulations of MuSCs are dissimilarly lost or retained in aging (Figures 1E-1F and 2A-2H). For example, rather than a stochastic loss, our scRNA-seq data shows that Cluster 1 (MuSC1) is more prone to age-related loss whereas the cells in Cluster 2 (MuSC2) display enhanced retention with age (Figure 1F). Importantly, this age-related shift in populations is more prominent in MuSCs compared to FAPs and macrophages, which seem to display a more stochastic change in populations during aging. Taking a closer look at MuSC population dynamics, MuSCs from young mice segregate into three major clusters based on UMAP clustering (Figures 2A-2B). MuSCs belonging to Cluster 3 (MuSC3) are characterized by genes involved in activation such as *MyoD1* and *Cdkn1c* (Figures 2B-2F and S1J). This cluster is completely lost in aged mice (Figure 2G) likely due to their propensity for early activation and their entry into cell cycle (Kimmel et al., 2020). On the other hand, MuSC Cluster 2 which is retained and augmented in aged mice displays increased expression of stress response genes, such as *Mt1, Mt2* and *Gpx3* (Figure S3). For example, Gpx3 has been shown to be required for the maintenance of the human MuSC pool (El Haddad et al., 2012). The activation of the stress response pathway by MuSC2 may provide these cells with resilience to cope with increased oxidative stress and genotoxicity of the aging niche (Figure S3).

**Figure 2.**
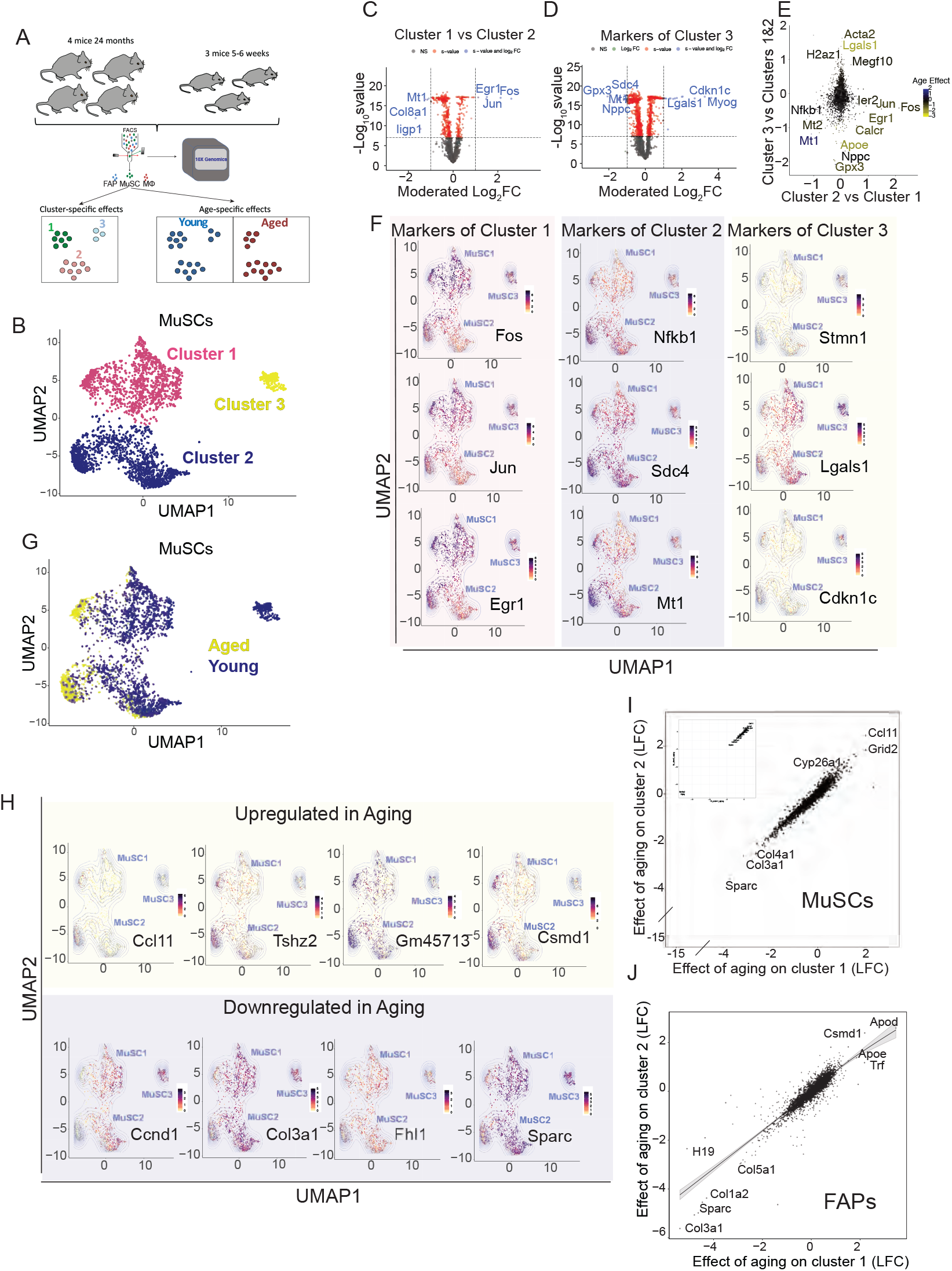
Aging Affects Common Genes Across Distinct Clusters of MuSCs. **(A)** Schematic diagram of the scRNA-seq analysis on young and aged MuSCs. **(B)** UMAP embedding of MuSCs, colored based on cluster. **(C)** Volcano plots showing genes that are up or downregulated in cluster1 compared to cluster2 of MuSCs, at a threshold of moderated LFC >1 and s-value <10^−8^. **(D)** Volcano plots showing genes that are up or down-regulated in cluster3 compared to Cluster1 & 2, at a threshold of moderated LFC >1 and s-value <10^−8^. **(E)** Scatterplot showing the moderated LFC of differentially expressed genes between Clusters MuSC2 and MuSC1, and between MuSC3 Cluster compared to Clusters MuSC1 & MuSC2. Blue and yellow represent down and upregulated genes respectively during aging. **(F)** Gene expression plots in the UMAP embedding for select representative positive marker genes of each cluster ranging from highest (black) to lowest (yellow) expression. **(G)** UMAP embedding of MuSCs, colored based on age. **(H)** Gene expression plots in the UMAP embedding for select representative genes that are up or downregulated in aging, ranging from highest (black) to lowest (yellow) expression. **(I)** Scatterplot showing the concordance of the moderated LFC effect of age on Clusters MuSC2 and MuSC1. **(J)** Scatterplot showing the concordance of the moderated LFC effects of age on FAP1 and FAP2 clusters in FAPs.

Since both MuSCs and FAPs display significant heterogeneity as evidenced by the presence of distinct clusters, we performed cluster-specific gene expression analysis between young and aged cells. Notably, we observe a strong correlation between age-affected genes on both of the main clusters of MuSCs, implying a consistent transcriptomic shift in different MuSC clusters that persist through aging (Figures 2H-2I). A similar effect is observed in FAPs (Figure 2J), which are present within the same niche. This data suggests that, despite differences in subpopulation loss, aging impinges upon similar gene networks and pathways, thus displaying an overarching effect.

### Heterochronic Transplantation Into a Young Niche Can Transcriptionally Reprogram Aged MuSCs

Given that aging affects similar gene networks and pathways regardless of the source subpopulation (Figures 2I-2J), in both MuSCs and FAPs, we reasoned that these alterations in the MuSC transcriptome may be directly caused by their niche environment during aging. We therefore proceeded to decouple age-related alterations in the MuSC transcriptome that are cell-intrinsic from changes caused by the aging niche. To assess this, we developed an *in vivo* model to directly quantify the effect of the niche on the MuSC transcriptome (Figure 3A). We first used FACS to isolate Pax7-nGFP MuSCs from the hindlimbs of mice aged 23-26 months (Figure 3A), followed by immediate injection of 10,000-20,000 of these freshly sorted aged donor MuSCs into the irradiated hindlimb of living young NOD-*Prkdc*^*em26Cd52*^*Il2rg*^*em26Cd22*^/NjuCrl (NCG) mice (Figures 3A-3D and Materials and Methods). Aged donor MuSCs were left to home to the young host niche for three weeks (Figure 3E). After 21 days of residing in the young niche, the engrafted Pax7-nGFP MuSCs were re-isolated by FACS (Figure 3F, Materials and Methods) for RNA-seq library preparation, to allow a direct transcriptome comparison of the same MuSCs before and after transplantation using SMART-Seq technology. To control for the computational modeling of the effect of engraftment itself, MuSCs from young mice (5-6 weeks old) were injected into host mice of similar age (Figure 3G, Materials and Methods). Since MuSC recovery yielded 30-400 cells on average, we used SMART-seq technology to create sequencing-ready libraries of cells before (T_0_) and after engraftment (T_21_), as it allows the reverse transcription of minute starting quantities of RNA (Picelli et al., 2014b) (Figure S4A-S4F, Materials and Methods). Genes altered by aging in our SMART-seq bulk libraries showed high concordance with age-affected genes from our independent scRNA-seq dataset (Figure S4G). After sequencing, genes altered between young and aged samples were classified into those showing an effect of age (different between young T_0_ and aged T_0_), an effect of engraftment/control (different between young T_0_ and young T_21_), and an effect of exposure to the heterochronic niche, defined as the effect of heterochronic transplantation while accounting for the effect of engraftment (Δniche=Δ_AgedT21-AgedT0_ – Δ_YoungT21-YoungT0_) (Figures 3G, 4A-4O). For example, age-affected genes that remain unchanged by the heterochronic niche include *B2m, Igf1, Cdkn3, Mt1*, and *Birc5*, amongst others, indicating the presence of cell-intrinsic age-related defects (Figures 4A-4C).

**Figure 3.**
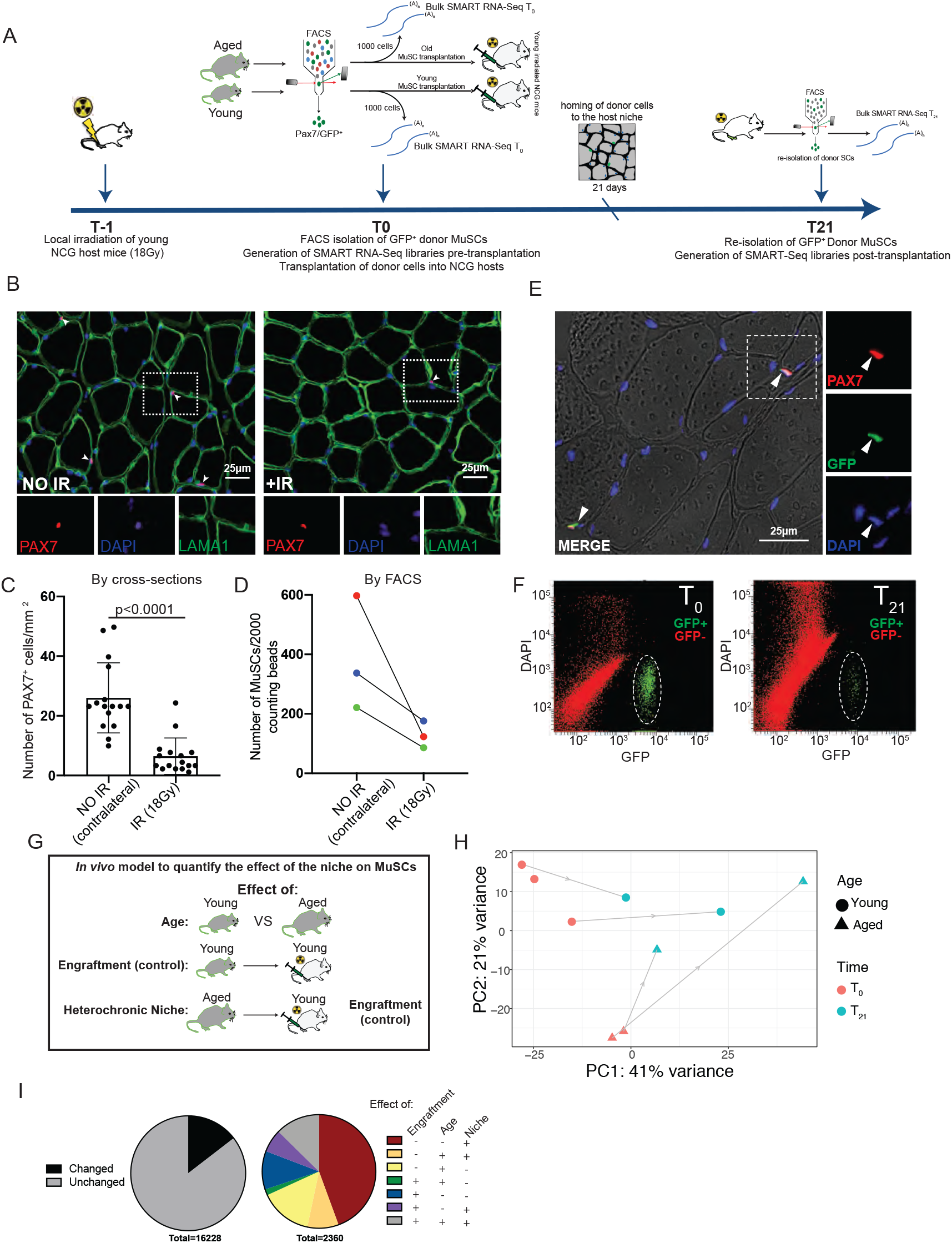
Development of an Allogeneic Stem Cell Transplantation Model to Directly Quantify the Effect of the Niche Environment on the Transcriptome of MuSCs. **(A)** Schematic diagram of the pipeline for allogeneic MuSC transplantation. **(B)** Immunofluorescent staining for PAX7, LAMA1 and DAPI on cross-sections from TA muscles that have been treated with 18Gy of irradiation compared to the contralateral untreated TA muscle (n=2 mice) **(C)** Quantification of the number of Pax7^+^ MuSCs/mm^2^ in irradiated compared to untreated contralateral Tibialis Anterior (TA) muscle cross-sections. (n=2 mice) **(D)** Quantification of the number of ITGA7^+^/Lin^-^ MuSCs per 2000 FACS counting beads sorted from irradiated compared to untreated contralateral TA muscles (n=3 mice) **(E)** Immunofluorescent staining for PAX7, GFP and DAPI of TA muscles from host mice showing engraftment and homing to the niche of GFP^+^ donor MuSCs 21 days post-transplantation. **(F)** Representative FACS plots showing the isolation of GFP^+^ MuSCs at T_0_ (left), followed by the re-isolation of the same cells after engraftment into young NCG mice at T_21_ (right). **(G)** Schematic diagram of the separation of age, engraftment, and niche effects after allogeneic MuSC transplantation. **(H)** Principal Component Analysis (PCA) plots of bulk RNA-seq data, from young and aged mice, pre-(T_0_) and post-transplantation (T_21_). **(I)** Pie chart of the relative proportion of genes that are altered after transplantation into the host niche versus the genes that remain unchanged after transplantation (left). Pie chart of the relative proportions of genes showing an effect of engraftment, age, or the allogeneic niche, in all combinations, from transplantation RNA-seq datasets (right). (moderated LFC >1, s-value < 0.05)

**Figure 4.**
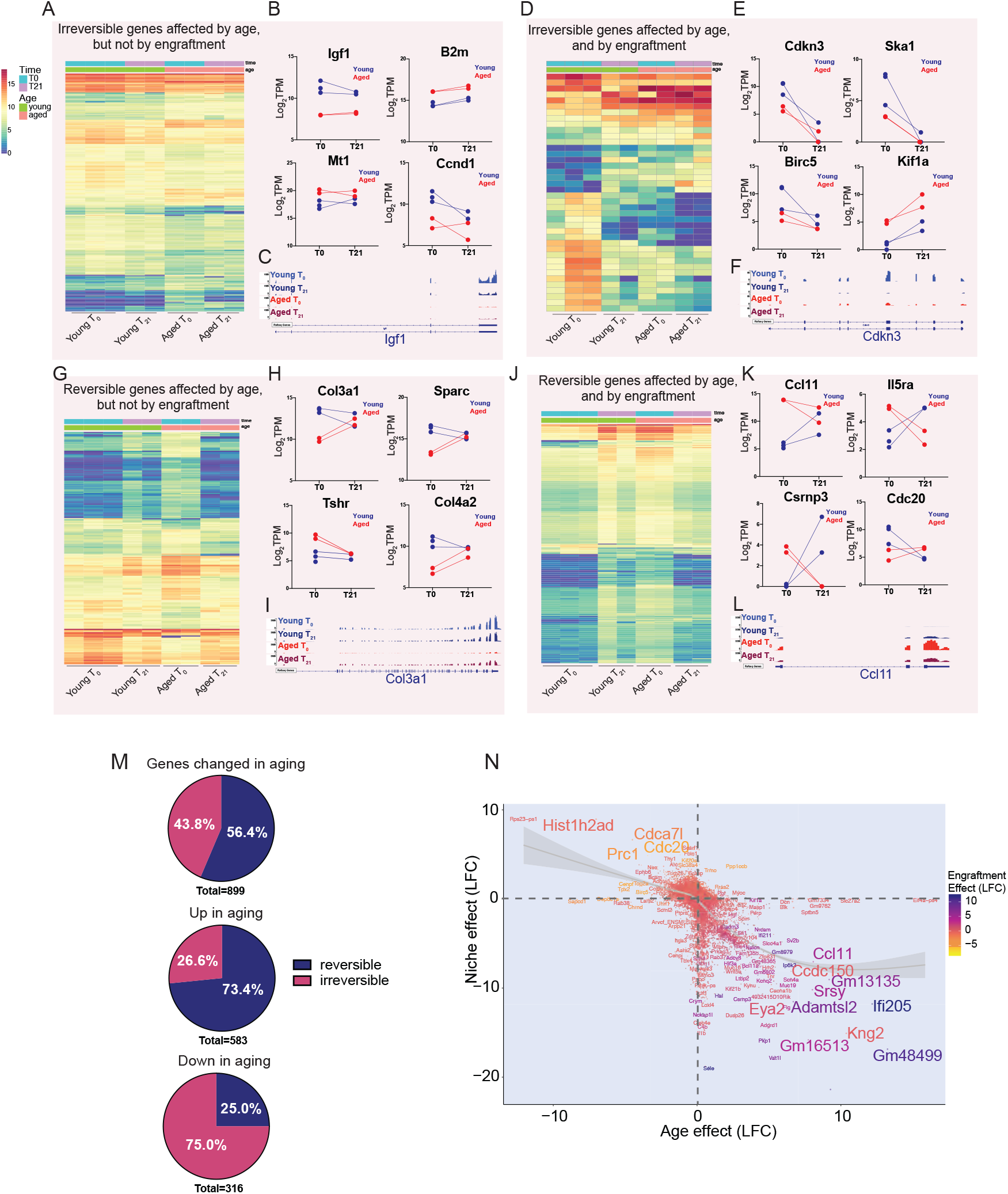
Exposure to the Young Niche Environment Restores a Significant Portion the Age-Related Altered Gene Expression Profile in MuSCs. **(A-C)** Heat maps (A) showing expression of genes affected by age, but not by engraftment or by the exposure to the niche, in hindlimb MuSCs (moderated LFC >1, s-value <0.05), followed by scatter plots (B) and Integrated Genome Viewer (IGV) tracks (C) for representative genes in this category. **(D-F)** Heat maps (D) showing expression of genes affected by age and engraftment, but not by niche exposure, in hindlimb MuSCs (moderated LFC >1, s-value <0.05), followed by scatter plots (E) and IGV tracks (F) for representative genes in this category. **(G-I)** Heat maps (G) showing expression of genes affected by age and niche exposure, but not by engraftment, in hindlimb MuSCs (moderated LFC >1, s-value <0.05), followed by scatter plots (H) and IGV tracks (I) for representative genes in this category. **(J-L)** Heat maps (J) showing expression of genes affected by age, engraftment, and niche exposure, in hindlimb MuSCs (moderated LFC >1, s-value <0.05), followed by scatter plots (K) and IGV tracks (L) for representative genes in this category. **(M)** Pie charts showing the percentage of genes that are up, downregulated, or changed in in either direction in aging that are either reversible or irreversible by exposure to the young niche (LFC >1, S-value <0.05). **(N)** Scatterplot of the moderated LFCs of the effect of niche exposure vs the effect of age.

To determine genes that are reversible by the niche environment back to youthful conditions after engraftment of aged MuSCs into young mice, we were interested in genes that show an effect of age and of the niche in their expression profile (Figures 4G-4L). At a threshold of s-value<0.05 and moderated LFC >1, out of 899 genes that are altered in aging, 507 are reprogrammable (56.4%) by their exposure to the young niche (Figure 4M). Notably, amongst these are genes involved in ECM composition and remodeling, such as *Col3a1, Col4a1, Col4a2, Mmp2*, and *Sparc* (Melouane et al., 2018), as well as known aging factor *Ccl11* (Villeda et al., 2011), and genes involved in cell cycle regulation, such as *Gas1* and *Cdc20* (Figures 4G-4L and S5). Interestingly, when plotting genes affected by age and by their response to heterochronic niche, a strong negative correlation between the two is immediately apparent, with a pronounced skew towards the restoration of genes that are originally upregulated in aging (Figure 5N). When quantifying this effect, we observed that 73.4% of genes that are up-regulated in aging are reversible, compared to only 25.0% of genes that are down-regulated in aging, at a threshold of moderated LFC >1 and an s-value <0.05 (Figure 4M). Overall, this data shows that the niche is a principal regulator of the MuSC transcriptome. More importantly, a significant portion of age-related altered transcriptome of MuSCs can be reprogrammed back to the youthful state by exposure to young niche milieu.

**Figure 5.**
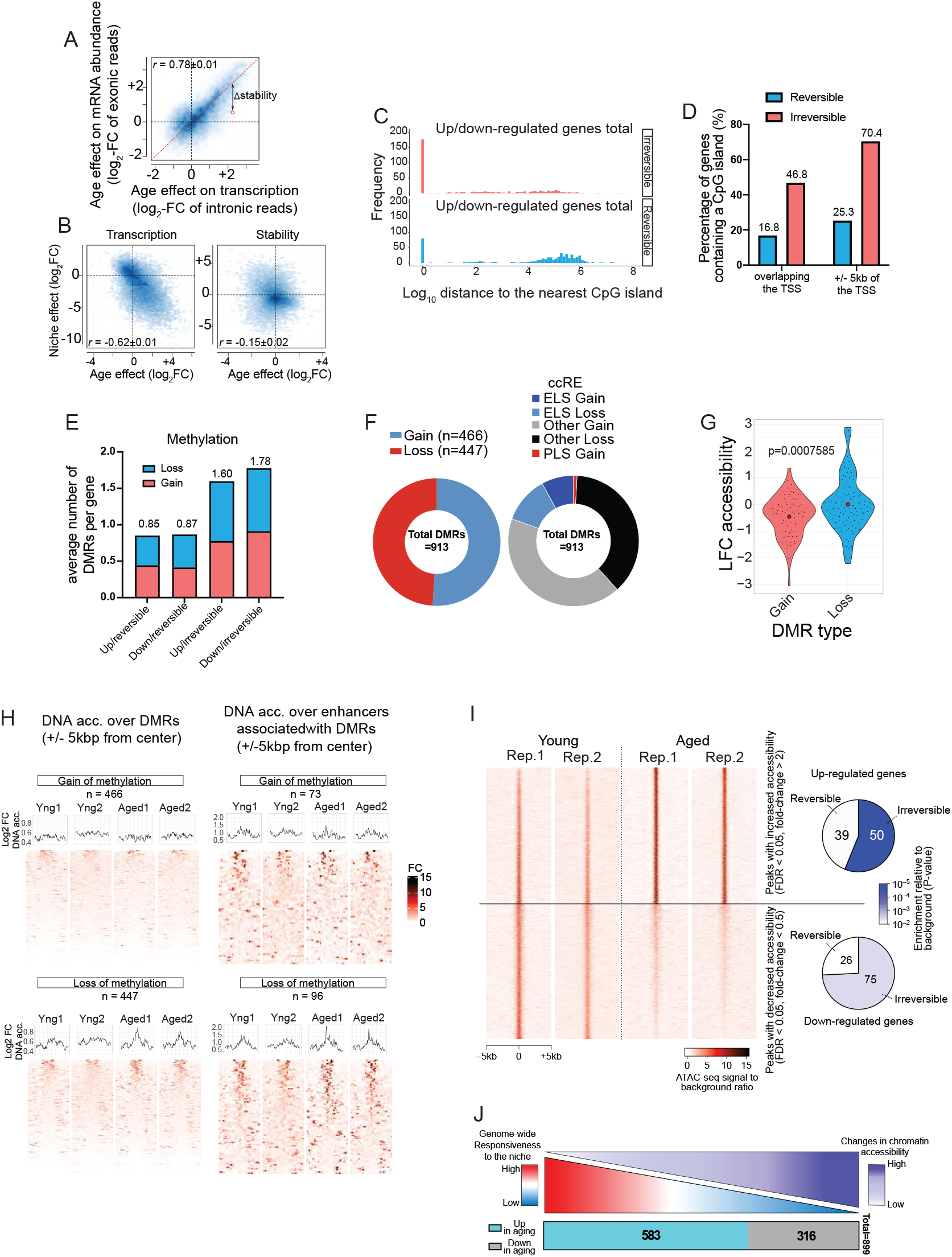
The Epigenetic State Dictates the Responsiveness of Age-Related Altered Genes to Heterochronic Niche Exposure. **(A)** Analysis of RNA stability. Scatterplot showing the age effect on transcription, as measured by changes in intronic read abundance between aged and young cells compared to the age effect of mature RNA abundance, based on exonic reads. Values along the diagonal represent genes that do not show significant difference in RNA stability between MuSCs from young and aged mice. **(B)** Scatterplots showing the effect of age compared to the effect of the heterochronic niche on transcription (left) and RNA stability (right). **(C)** Frequency of CpG islands within the vicinity of the TSS of genes that are up- or downregulated in aging. **(D)** Quantification of the percentage of reversible and irreversible genes containing a CpG island overlapping with or within 5kb of their TSS. **(E)** Quantification of the average number of DMRs associated with each gene category, as determined by the number of DMRs at cis-regulatory elements associated divided by the number genes in each category. **(F)** Pie-chart showing the proportion of age-related DMRs corresponding to a gain or loss of methylation (left).Pie Chart showing the proportion of identified age-related DMRs associated with cCREs (+/- 300bp) that are located within Enhancer-like sequences (ELS), promoter-like sequences (PLS), or other regions of the genome (right). **(G)** Violin plots showing the LFC in MuSC chromatin accessibility and age-related DMRs **(H)** ATAC-seq pileup analyses showing chromatin accessibility either over DMRs (+/- 2KB) or over enhancers associated with DMRs in young and aged MuSCs. **(I)** Left: Heatmap of differentially accessible ATAC-seq peaks, including peaks with increased accessibility in aged cells (up) and those with decreased accessibility (bottom). Right: Pie-charts showing the overlap of differentially accessible peaks with different gene categories. The top pie chart shows the number of reversible/irreversible age-up-regulated genes that are associated with at least one age-up-regulated peak. The bottom pie chart shows the number of reversible/irreversible age-down-regulated genes that are associated with at least one age-down-regulated peak. The color gradient corresponds to the P-value of enrichment of each category, relative to the background frequency (Fisher’s exact test). **(J)** Schematic overview showing the relationship between the responsiveness to the niche of genes that are up- and down-regulated in aged MuSCs and alternations in chromatin accessibility.

### The Epigenetic Signature Dictates the Transcriptional Response of MuSCs to the Niche Environment

To gain mechanistic insight into the age-related alterations in the transcriptome of MuSCs and the effect of the niche environment on their transcriptional reprogramming, we first asked whether post-transcriptional and/or epigenetic mechanisms, such as RNA stability and chromatin state respectively, may be playing a role. We started by examining the RNA stability changes associated with aging and/or niche response in MuSCs using the RNA-seq data described in the previous section (Materials and Methods). We deconvolved the transcriptional and post-transcriptional effects based on the relative abundances of pre-mRNA and mature mRNA (Perron et al., 2021) (Materials and Methods), and determined that differential RNA stability is not a primary mechanism of age-related altered transcriptome and niche-mediated restoration of age-related genes (Figures 5A-5C). An alternate possibility is that transcriptional reprogramming occurs at the chromatin level. We therefore hypothesized that the previously-observed differential response between up versus downregulated genes may be a manifestation of different chromatin states (Figure 4N). To further delve into the differential reversibility of genes that are upregulated versus downregulated in aging, we assessed the frequency of CpG islands within the vicinity of the TSS of age-related altered genes as a proxy for the capacity to be influenced by DNA methylation (Figures 5C-5D). Interestingly, we observed that the highest frequency of CpG islands is associated with irreversible genes (i.e., genes not responsive to niche exposure) (Figures 5C-5D). Next, we performed Whole Genome Bisulfite Sequencing on primary myoblasts derived from young and aged MuSCs (Figure S7A-S7C, Materials and Methods). Interestingly, age-related irreversible genes in MuSCs are associated with the highest frequency of DMRs within candidate *cis*-regulatory elements (cCREs) (Figure 5E). We particularly observed a significant over-representation of irreversible age-upregulated genes among those associated with loss-of-methylation DMRs (odds ratio 2.2, P<0.05) (Table 1).

**Table 1.** Table showing the number of DMRs overlapping with genes that are upregulated in aging, downregulated in aging, reversible and irreversible by the niche.

|  |  |  | Groups of genes of interest |  |  |  |  |  |  |
| --- | --- | --- | --- | --- | --- | --- | --- | --- | --- |
|  |  |  | All groups | Downregulated |  |  | Upregulated |  |  |
|  |  |  |  | All downregulated | irreversible | reversible | All upregulated | irreversible | reversible |
| Genes associated to DMRs | Gain | p-value | 0.7638 | 0.693 | 0.4029 | 1 | 0.7358 | 0.8981 | 0.505 |
|  |  | odds ratio | 0.8486 | 0.8691 | 1.171 | 0 | 0.8362 | 0.5131 | 1.0626 |
|  |  | # of overlapping genes | 15 | 7 | 7 | 0 | 8 | 2 | 6 |
|  | Loss | p-value | 0.663 | 0.7426 | 0.6203 | 0.876 | 0.5383 | **0.042** | 0.9851 |
|  |  | odds ratio | 0.9206 | 0.8226 | 0.9372 | 0.4785 | 1.008 | **2.201** | 0.3171 |
|  |  | # of overlapping genes | 17 | 7 | 6 | 1 | 10 | **8** | 2 |

| All genes of interest by group (intersect with background) |  |  |  |  |
| --- | --- | --- | --- | --- |
| down_irrev | down_rev | up_irrev | up_rev | Total |
| 95 | 30 | 59 | 89 | 273 |

On a global scale, we observed no net gain or loss of methylation at DMRs, which is consistent with previous studies (Figure 5F) (Hernando-Herraez et al., 2019). Similar to what has been shown in various cancers (Angeloni and Bogdanovic, 2019; Blattler et al., 2014), most DMRs map outside of the Transcription Start Sites (TSS), indicating a potential role for enhancer elements (Figure 5F). Next, to determine if changes in DNA methylation lead to changes in chromatin accessibility, we performed Assay for Transposase-Accessible Chromatin using sequencing (ATAC-seq) on 5000 freshly sorted MuSCs isolated from young and aged mice to compare the status of chromatin accessibility of age-deregulated genes (Figures S6, S7D-S7F) and Materials and Methods). By assessing chromatin accessibility spanning the DMRs, we observe that age-related loss of methylation leads to an increase in chromatin accessibility whereas gain in methylation leads to a decrease in accessibility (Figures 5G-5H). In fact, we identified a total of 7258 ATAC-seq peaks with significant age-associated changes in chromatin accessibility (LFC >1, p_adj_ <0.05). Importantly, genes that are irreversible are more likely to be associated with a change in chromatin accessibility compared to the reversible counterparts (Figure 5I and Table 2). Specifically, we observed significant overlap between irreversible up-regulated genes and those associated with age-accessible cCREs (P< 5×10^−6^), and irreversible down-regulated genes and those associated with age-inaccessible cCREs (P< 0.004) (Figure 5I). Together, this data suggests that irreversible age-related changes in the transcriptome are driven by the chromatin state in cCREs, and that distinctive epigenetic signatures can influence the responsiveness to niche-induced transcriptional reprogramming (Figure 5J).

**Table 2.** Table showing the number of ATAC-seq peaks overlapping with genes that are upregulated in aging, downregulated in aging, reversible and irreversible by the niche.

|  |  |  | Groups of genes of interest |  |  |  |  |  |  |
| --- | --- | --- | --- | --- | --- | --- | --- | --- | --- |
|  |  |  | All groups | Downregulated |  |  | Upregulated |  |  |
|  |  |  |  | All downregulated | irreversible | reversible | All upregulated | irreversible | reversible |
| Genes associated to ATAC-seq peaks | Upregulated peaks | p-value | 0.00107 | 0.5016 | 0.5412 | 0.4764 | 3.2127e-06 | 9.8406e-07 | 0.067 |
|  |  | odds ratio | 1.3272 | 1.0064 | 0.9947 | 1.0441 | 1.8621 | 2.5469 | 1.3571 |
|  |  | # of overlapping genes | 245 | 119 | 90 | 29 | 126 | 72 | 54 |
|  | Downregulated peaks | p-value | 6e-04 | 2.6617e-07 | 0.00035 | 2.4459e-05 | 0.8135 | 0.9653 | 0.3198 |
|  |  | odds ratio | 1.3467 | 1.8456 | 1.6081 | 2.8869 | 0.8935 | 0.7161 | 1.1137 |
|  |  | # of overlapping genes | 267 | 172 | 124 | 48 | 95 | 42 | 53 |

| All genes of interest by group (intersect with background) |  |  |  |  |
| --- | --- | --- | --- | --- |
| down_irrev | down_rev | up_irrev | up_rev | Total |
| 217 | 68 | 112 | 110 | 507 |

## Discussion

The extent to which the age-related changes in MuSC function are a cause or consequence of aging muscle tissue has important implications for our understanding of skeletal muscle regeneration in healthy and aging skeletal muscle. Here, we decouple age-related cell-intrinsic effects, niche-mediated cell-extrinsic effects, and changes in population dynamics of MuSCs and two key muscle-resident cells in young and aged mice. First, we report that the age-related reduction in MuSCs is not stochastic, as specific subpopulations are preferentially lost and or retained in aged mice (Figures 1 and 2). The differential loss/retention of subpopulations of MuSCs such as MuSC2 may be in part due to their ability to upregulate stress response genes, giving them protection against a genotoxic environment during aging.

Remarkably, the effect of age is primarily uniform across different MuSC clusters despite major differences in their transcriptome as demonstrated by scRNA-seq. The uniform effect of age on the alteration of common genes and pathways in both MuSCs and FAPs suggests that the niche environment plays a fundamental role in the alteration of gene expression associated with age (Figure 2I). While it is known that aging affects the composition of the MuSC niche, the direct effect of the niche on the gene expression of the residing stem cells has remained unexplored. Our allogeneic transplantation model (Figure 3) shows that a significant portion of age-related changes in the transcriptome can be reversed by exposure to the young muscle environment, suggesting that the changing niche is a principal driver of the altered MuSC transcriptome (Figure 4). Therefore, as opposed to focusing solely on MuSCs themselves, bioengineering of the niche in its entirety presents a viable therapeutic approach to mitigate the effects of aging on MuSCs.

Importantly, despite the high percentage of gene reversibility observed, our allogeneic transplantation model allowed us to tease out genes that are irresponsive to the niche. Our data shows that the age-deregulated alterations that are encoded epigenetically are less responsive to the niche compared to changes in the transcriptome that are not associated with altered chromatin accessibility or DMRs (Figure 5). More specifically, our data shows that genes that are irresponsive to the young niche are associated with age-related changes in chromatin and in DNA methylation. On the other hand, genes that can be reversed are not linked to changes in chromatin or epigenetic signature during aging. This concept is consistent with what has been previously shown in the field of stem cell reprogramming, where the remodeling of cells’ epigenetic signature by forced expression of transcription factors is needed to return cells to a more pluripotent or progenitor-like state (Ocampo et al., 2016; Takahashi and Yamanaka, 2006). Similarly, our data shows that the transcriptional response of MuSCs to the niche environment is dependent on their epigenetic signature, suggesting that alterations in chromatin accessibility and DNA methylation impede transcriptional restoration by niche exposure.

Overall, our data highlights the importance of the muscle stem cell niche environment as a critical regulator of MuSC gene expression. More importantly, the significant plasticity in the restoration of the age-related altered transcriptome of MuSCs suggests that modulating the niche environment can be a powerful tool to restore stem cell-mediated endogenous muscle regeneration in aging. Our data also has important implications with regards to preventative therapies, suggesting that any potential treatments aiming to target the niche would be most effective prior to both the initiation of age-related changes in population dynamics and the onset of alterations in the chromatin state.

## Materials & Methods

### Isolation of Pure Populations of MuSCs and Niche Cells by Fluorescence-Activated Cell Sorting (FACS)

Hindlimb muscles were dissected from young (4-6 weeks old) or aged (23-26 months old) mice and minced until no visible tissue chunks were visible. Muscle was then digested in a 15 ml Falcon tube containing 2.4 U/ml Collagenase D (11088882001, Roche), 12 U/ml Dispase II (04942078001, Roche) and 0.5 mM CaCl_2_ in Ham’s F10 media (11550043, Gibco) for two periods of 30 minutes each on a shaker in a 37 °C/5.0% CO_2_ Tissue Culture (TC) incubator. After the first 30 minutes, the remaining muscle chunks were pelleted by centrifugation at 600 xg for 25 seconds and the supernatant containing released mononuclear cells was supplemented with 9 ml Fetal Bovine Serum (FBS) (080450, Wisent). The remaining pellet of undigested muscle tissue was triturated, and additional digestion buffer was added for the second 30-minute incubation. The combined digested tissue was then filtered through a 40μm cell strainer (C352340, Falcon) and spun down at 600 xg for 18 minutes. The supernatant was discarded and the cell pellet was dissolved in 500 μl 2% FBS/PBS (v:v) with 0.5 mM EDTA, and filtered once more. For isolation of GFP+ MuSCs from Pax7-nGFP mice, 0.5 μg of DAPI (Invitrogen, D3671) was added for cell sorting of the DAPI^-^/GFP^+^ population using a FACSAria Fusion cytometer (BD Biosciences). For simultaneous isolation of MuSCs, FAPs and macrophages, the digested tissue was supplemented with FC (fragment crystallizable) block (clone 2.4G2). Antibodies for anti-mouse ITGA7-AF647 (R&D systems, FAB3518R), anti-mouse CD31-PE (Invitrogen 12-0311-81), anti-mouse Sca1-FITC (BD Pharmigen, 557405), anti-mouse CD45-BV785 (Biolegend, 103149), anti-mouse CD11b-PerCP-Cy5.5 (Biolegend, 101227), anti-mouse F4/80-APC-Cy7 (Biolegend, 123117), and Hoechst 33342 (H1399, Molecular Probes) were added and the sample was incubated at room temperature (RT) for 15 minutes with intermittent shaking. The sample was then washed with 5 ml of 2% FBS/PBS (v:v) with 0.5 mM EDTA and spun down at 600 xg for 15 minutes. The stained cells were then resuspended in 700 μl of 5% FBS/PBS (v:v) with 0.5 mM EDTA and filtered through a 40 μm filter before sorting. Color compensation was done by staining 20 μl of UltraComp eCOMP beads (01-2222-41, Invitrogen) in 200 μl of 2% FBS/PBS (v:v) and calculated using DIVA before sorting on a FACSAria fusion III cytometer (BD Biosciences). In order to maintain the ratios of MuSCs, macrophages and FAPs, all cells were sorted into a single tube and 2500 cells were captured per sample for single-cell RNA-Sequencing using 10X Genomics chromium platform. FACS gating strategies benefited from *Pasut A, et al*, 2012(Pasut et al., 2012), *Farup J, et al*, 2015(Farup et al., 2015), and *Low et al*, 2017(Low et al., 2017).

### Allogeneic MuSC Transplantation into a Heterologous Niche

NOD-*Prkdc*^*em26Cd52*^*Il2rg*^*em26Cd22*^/NjuCrl (NCG) immunocompromised mice (Charles-River) were irradiated on their hindlimb the day before transplantation. Mice were anesthetized by IP injection of a rodent cocktail composed of ketamine (100 mg/kg)/xylazine (10 mg/kg)/acepromazine (3 mg/kg) and their hindlimb was positioned into the field of irradiation of a Multirad225 irradiation machine (Precision X-Ray) and were given a 18Gy dose of irradiation as described previously (Morgan et al., 1990; Morgan et al., 1993; Zismanov et al., 2016). The next day, MuSCs were isolated from Tg(Pax7-EGFP)#Tajb(Sambasivan et al., 2009) mice as described previously, and sorted into a 2% FBS/PBS mix. 10 000-20 000 freshly-isolated GFP^+^ donor MuSCs were immediately injected into the Tibialis Anterior (TA) muscle of the previously irradiated hindlimb of host NCG mice using a Hamilton Syringe, in 2-3 separate injection sites along the TA muscle.

### Immunostaining of TA Muscles

TA muscles were dissected tendon to tendon and rinsed in cold PBS. The isolated muscles were then fixed in 0.5% Paraformaldehyde in PBS for 2.5 hours at 4 °C and then placed in 20% Sucrose (S7903, Sigma) at 4 °C overnight. Following 12-24 hours, muscle samples were frozen in aluminum foil cups of Optimal Cutting Temperature (OCT) using liquid nitrogen-chilled isopentane and stored at −80 °C prior to sectioning by the cryostat at 8 μm interval. TA muscle cross-sections on glass slides were circled using a hydrophobic PAP pen. Cross-sections were permeabilized in 0.2% Triton X-100/0.1M glycine in PBS for 7 minutes on a shaker, and then washed in PBS. Cross-sections were blocked in 3% (m/v) BSA/10 % Goat Serum solution for at least 1 hour at RT in a humid chamber. TA muscle sections were then incubated with primary antibodies for PAX7 (DSHB, AB_528428), GFP (Invitrogen, A-11120), and/or LAMA1 (Sigma, L9393) diluted in blocking solution in a humid chamber at 4 °C overnight, then washed 3 times for 10 minutes with PBT (0.05% Triton X-100 in PBS). Secondary antibodies diluted in blocking solution were added for one hour at RT in a humid chamber, followed by 3 × 10 minute washes with PBT. ProLong Gold Antifade Solution with DAPI (P3695, Invitrogen) was used for mounting prior to imaging.

### Analysis of Cell Purity Post FACS Isolation

Cells were freshly sorted by FACS into a 1.5 ml Eppendorf tube containing 100 μl of 2% FBS/PBS (v:v). Freshly sorted cells were pelleted by centrifugation at 600 xg and resuspended in 50 μl of PBS. The cells were then plated in a well of a 12-well plate, and excess liquid was allowed to evaporate for around 10 minutes, or until wells appeared dry. Cells were then fixed using 3.2% PFA/PBS to the well, gently to not dislodge the cells, for 10 minutes at RT. After washing 3x with PBS, cells were permeabilized with 0.5% Triton X-100 for 30 minutes at RT. Cells were then washed 2x with 0.3% Triton X-100/PBS and blocked with 0.3%Triton X-100/0.5% BSA/PBS for 1 hour at RT. Primary antibodies for PDGFRA (Cell signaling, 31745), F4/80 (Invitrogen, 124801-82), and PAX7 (DSHB, AB_528428) were diluted in blocking solution and added for an overnight incubation at 4 °C in a humid chamber. Next, the cells were washed 3x with 0.3% Triton X-100 before the addition of the secondary antibodies, diluted in blocking buffer for 1 hour at RT. Finally, cells were washed 2x with 0.3% Triton X-100 before the addition of PBS with DAPI and visualized on a microscope.

### Single-cell RNA-Sequencing (scRNA-seq)

MuSCs, macrophages and FAPs were isolated from three young (5 weeks old) and four aged (23.5 months old) mice by FACS, as described previously, and immediately processed for single-cell RNA-sequencing (scRNA-seq) using Chromium Single Cell 3’ Reagent Kits (v3 Chemistry) using 10x genomics technology. In order to maintain ratios, all cells were sorted into a single tube and to aim for a total of 2500 captured cells for sequencing per biological replicate. The viability of cells was 85.5% on average, with a capture range of 2437 to 2835 cells. Quality control and processing steps can be found at https://csglab.github.io/aging_muscle_niche/pages/notebooks.html).

### SMART-seq Library Preparation

MuSCs were sorted by FACS directly into 9 μl of nuclease-free water containing 1 μl SMART-Seq Reaction Buffer, which is composed of 10x lysis Buffer supplemented with RNase inhibitors. The volume was brought to 11.5 μl and libraries were processed using the SMART-Seq HT library preparation kit (634456, Takara Bio) as described (Blackburn et al., 2019; Picelli et al., 2014b). cDNA libraries were then purified using Ampure XP beads (A63880, Beckman Coulter) at a 1:1 (v:v) ratio and quantified using the Quant-iT PicoGreen dsDNA Assay kit (P11496, Invitrogen). 0.15 ng of cDNA in a 1.25 μl volume was used as input for Nextera XT (FC-131-1024, FC-131-2001, Illumina) tagmentation, as described (Adey et al., 2010; Picelli et al., 2014a). Sequencing libraries underwent Ampure XP size selection using a 1:0.85 (v:v) ratio, and an aliquot was used to verify size on an agarose gel stained with GelGreen dye (41005, Biotium). Final libraries underwent quality control verifications using a bioanalyzer and were sequenced on an Illumina NextSeq500.

### Assay for Transposase-Accessible Chromatin Using Sequencing (ATAC-seq)

Low input ATAC-seq (Assay for Transposase-Accessible Chromatin using sequencing) was performed according to the OMNI ATAC-seq (Corces et al., 2017) protocol. Briefly, 5000 MuSCs were FACS sorted into the lysis buffer (10 mM Tris-HCl pH 7.5, 10 mM NaCl, 3 mM MgCl_2_, 0.1% Tween-20, 0.1% NP-40, 0.01% Digitonin) and incubated for 5 minutes on ice and subsequently 3 minutes at room temperature (RT). Cells were then washed with 100 μl of wash buffer (10 mM Tris-HCl pH 7.5, 10 Mm NaCl, 3 mM MgCl_2_, 0.1% Tween-20) and spun for 800g for 10 minutes. The pellet was resuspended in the transposition mixture at a total volume of 10 μl (5 μl of Tagment DNA (TD) buffer, 3.2 μl PBS, 0.89 μl Tn5, 0.1% Tween-20, 0.01% Digitonin and 0.75 μl water). Transposition was performed for 20 minutes at 37 °C while shaking the tubes every 5-7 minutes. The DNA was then purified using column purification as described (Qiagen, QIAquick PCR Purification Kit Cat: 28104). The purified DNA was then PCR-amplified using Q5 DNA polymerase for 12 cycles with the Illumina Nextera XT adapters. The amplified DNA was then size selected and purified with Ampure XP beads at a 1:0.85 (v:v) ratio. The quality control of the final libraries were performed by bioanalyzer and paired-end sequencing was performed on NovaSeq 6000 Sprime paired end (PE 150 bp).

### Genomic DNA Extraction

Genomic DNA was extracted using phenol/chloroform/isoamyl alcohol solution (Invitrogen #15593-031). Briefly, primary myoblasts derived from *in vitro* expansion of freshly sorted MuSCs were pelleted at 600 xg for 10 minutes and snap frozen in liquid nitrogen before use. When all samples were collected, cell pellets were thawed on ice and lysed in 400 μl lysis buffer (40 nM Tris-HCl pH 8.0; 1% (vol/vol) Triton X-100; 0.1% (vol/vol) SDS; 4 mM EDTA (pH 8.0) and 300 mM NaCl). Cell lysates were incubated at 37 °C for 45 minutes with 10 μl RNase A (Invitrogen, cat#12091-021), followed by a 1h 15min incubation at 45 °C with 10 μl of Proteinase K (10mg/ml Millipore, cat#70663). One volume of phenol/chloroform was then added to each lysate, and samples were vortexed vigorously and centrifuged at 2110 xg for 3 minutes. The aqueous layer was retained and mixed with 2 volumes of 100% ethanol (EtOH). Samples were snap frozen in liquid nitrogen for 3 minutes, thawed on ice, and centrifuged for 20 minutes at 2110 xg. The DNA pellet was washed 3x with 750 μl of 75% EtOH, spinning each time 2110 xg for 12 minutes. The DNA pellet was then air-dried and resuspended in TE buffer.

### Whole-Genome Bisulfite Sequencing (WGBS) Library Preparation

Genomic DNA was sheared using COVARIS, and libraries were constructed using the NxSeq AmpFREE low DNA Library Kit by Lucigen. Bisulfite conversion was done using the EZ-96 DNA Methylation-GoldTM MagPrep kit (D5042, Zymo Research).

### Analysis of Differentially Methylated Regions (DMRs) after Whole Genome Bisulfate Sequencing

Quality check and adaptor trimming of raw reads was performed by trim-galore (Default setting, Phred score = 20) (Martin, 2011). Reads were aligned to the mouse reference genome (mm10) using bismark (internal bowtie2 aligner’s default settings and allowing one mismatch) (Langmead and Salzberg, 2012). Extraction of methylation calls was done by ignoring the first 2 base pairs from the 5’ end of read 2 to avoid known experimental-introduced bias (Krueger and Andrews, 2011).

The DMRs between young and aged samples, were identified with DMRcaller’s (R/Bioconductor) function computeDMRsReplicates (Catoni et al., 2018). DMRcaller implements a Beta regression to test the difference in methylation levels between conditions in regions of 100 bps with at least 4 CpG sites. Only regions with a minimum methylation proportion difference of 40% are reported. P-values are adjusted for multiple testing using Benjamini and Hochberg’s method (adjusted p-value < 0.01) (Benjamini and Hochberg, 1995).

### Bulk RNA-seq Data Analysis

Transcript-level abundances were imported into R and collapsed into gene counts using the *tximport* package (Soneson et al., 2015). To estimate the effects of the experimental conditions, we used *DESeq2 (Love et al., 2014)* to fit the following gene-wise model: *count∼ time + age + time:age*, where the variable *time* denotes the effect of engraftment, the variable *age* denotes the effect of aging, and the interaction variable *time:age* represents the reprogramming effect on genes in the aged niche (i.e. the residual effect beyond that of engraftment or aging that explains observed expression of a given gene). For the purposes of ranking and visualization, we used the *apeglm* shrinkage approach (Zhu et al., 2019).

### Single Cell RNA-seq Data Analysis

We used *Salmon* and a*levin* (Patro et al., 2017; Srivastava et al., 2019) to quantify spliced mRNA and unspliced pre-mRNA abundances from scRNA-seq data. To create a spliced-intronic Salmon quantification index, we used *eisaR* (Soneson et al., 2021) to generate a combined FASTA file containing exonic and intronic sequences from the mouse Genocode M24 transcriptome and genome reference files. The introns were defined using the “collapse” option in which transcripts of the same gene are first collapsed by taking the union of all the exonic regions within a gene and labeling the remaining non-exonic parts as introns. In contrast to the alternative “separate” strategy in which intronic regions are extracted for each transcript separately, the collapse strategy treats an ambiguous read to be more likely to have come from an exon of a given transcript than from an intron of alternate transcript that happens to overlap that exonic region (Soneson et al., 2021). A 90 nt flanking sequence was added to each side of each extracted intronic region (equal to the length of Read 2 in Chromium Single Cell 3’ Gene Expression library with v3 chemistry) in order to quantify intron-exon reads as unspliced pre-mRNA abundances. To exclude reads coming from intergenic regions of the genome, we adopted the selective alignment (SA) strategy by providing Salmon the complete genome sequence as a decoy sequence (Srivastava et al., 2020).

The alevin quantifications were imported into *R 4*.*0*.*0* using the *tximeta* package. The abundances were split into a spliced and unspliced abundances matrices to be used for downstream analysis such as RNA velocity. We identified and removed low-quality or potentially multiplet cells based on whether they had high percentage of mitochondrial reads (>15%), low library sizes, low number of genes detected, or whether they tend to be observed as outliers with respect to the distributions of spliced reads, unspliced reads, or the fraction of unspliced reads in each cell (see the analysis notebooks for details: https://csglab.github.io/aging_muscle_niche/pages/notebooks.html).

To cluster cells, we used UMAP (McInnes et al., 2018) followed by the Walktrap community finding algorithm (Pons and Latapy, 2005) to reduce the dimensionality of the data and pull together similar cell populations into separated clusters which can then be labelled. To annotate the main clusters, we used the expression of *Adgre1, Ptprc*, and *Itgam* as markers of macrophages, *Pax7* as a marker of muscle stem cells (MuSCs), and *Pdgfra* as a marker of FAPs. The initial clustering showed the presence of three large main clusters corresponding to the three cell types along with four small populations whose expression profiles suggest them as being Schwan and smooth muscle cell clusters respectively (Figure S1E-S1G). The four small cell clusters were ignored in downstream analysis. Next, the seven samples were batch normalized and corrected for batch effects using fast mutual nearest neighbors (MNN) correction (Haghverdi et al., 2018). The clustering approach was repeated independently on the MNN corrected space of the MuSCs, FAP, and macrophages respectively in order to determine the internal heterogeneity of each cell type cluster. As a result, MuSCs were partitioned into three clusters, the FAPs into two clusters, and the macrophage cells into one cluster only.

We ran differential gene expression analysis separately for each cell type. We discarded genes that had low counts, that had low mean expression across the subclusters making up the cells of each cell type, and that were characterized by severe batch effects. We used ZINB-WaVE (Risso et al., 2018) to compute gene and cell-specific observational weights that can be used to unlock *DESeq2 (Love et al., 2014)* for the differential analysis of single cell RNA-seq data. For data of each of the three cell types, we independently fit the following gene-wise model: *count∼* −1 + *cluster_condition + sample*, where the variable *cluster_condition* denotes the mean expression of cells in each cluster and age condition, and the variable *sample* denotes the batch effect of each sample. We manually created the design matrix such that the sample effects of one of the aged samples and one of the young samples are removed in order to fit a statistical identifiable model. We tested several contrasts to assess several hypotheses of differential mean gene expression across the different sub clusters. For the purposes of ranking and visualization, we used the *ashr* shrinkage approach (Stephens, 2017) to preserve the size of estimated large LFC and compute *s-values* (the estimated rate of false sign).

### Integration of Bulk and Single-cell RNA-seq Data

Using the moderated LFCs generated by models fit on both RNA-seq data modalities, we are able to examine the pattern of concordance of differential gene expression across the multiple datasets. More specifically, the moderated LFC model effect estimates for the bulk data and the muscle stem cells (MuSCs) single cell data were joined together such that we retain only genes that were not discarded from the scRNA-seq data. The LFCs were used for plotting Figure S4G.

### Geneset Enrichment Analysis

We used the R package hypeR (Federico and Monti, 2020) to conduct hypergeometric tests to determine which respective set of statistically significant differentially regulated genes are over-represented in the 50 pre-defined hallmark gene sets listed in the Molecular Signatures Database (MSigDB)(Liberzon et al., 2015). The threshold for significance was set at 10-8 for the s-values across the contrast effects accompanied with an absolute value cutoff for the moderated LFCs between 0.5 and 2 depending on the distribution of effect values for each contrast. The six sets of significant genes for MuSCs are the following: (1) genes that are downregulated in aging both in bulk and single cell, (2) genes that are downregulated in aging both in bulk and single cell (scRNA-seq), (3) genes that are upregulated in MuSC2 with respect to MuSC1 cluster, (4) genes that are upregulated in MuSC1 with respect to MuSC2 cluster, (5) genes that are upregulated in MuSC3 with respect to young cells in both MuSC1 and MuSC2 clusters cluster, (6) genes that are downregulated in MuSC3 with respect to young cells in both MuSC1 and MuSC2 cluster. For FAPs, there were four sets: (1) genes that are downregulated in aging (2) genes that are upregulated in aging (3) genes that are upregulated in FAP 2 with respect to FAP1, (4) genes that are upregulated in FAP1 with respect to FAP2.

### ATAC-seq Pre-Processing Analysis

The ATAC-Seq data was processed using the ENCODE ATAC-seq pipeline (https://github.com/ENCODE-DCC/atac-seq-pipeline). The pipeline consists of adapter read trimming, read alignment, post-alignment read filtering and de-duplicating, calling peaks, creating signal tracks, generating quality control reports, and running Irreproducible Discovery Rate (IDR) analysis (Qunhua et al., 2011). The full specification of the pipeline is detailed on the website (https://github.com/ENCODE-DCC/atac-seq-pipeline). For downstream analysis the generated IDR optimal peaks were used.

### Analysis of Differentially Bound ATAC-seq Peaks

Peaks from young and aged conditions were merged from the IDR pipeline. Peaks with summits within 200 bp were combined, a new summit was set in the middle. The peaks’ start and end were redefined with a range of 500 bp from the summit.

We created a count matrix by extracting ATAC-seq read counts from peak and background (+/- 10kbp around the peak) regions for all four samples, peaks and samples correspond to rows and columns respectively (Quinlan, 2014). We used DiffRAC and the count matrix to identify significant differentially bounded peaks (adjusted p-value < 0.05 and LFC > 1) (Perron et al., 2021).

### Analysis of Differential mRNA Stability

Differential mRNA stability was inferred as previously described (Perron et al., 2021). Briefly, RNA-seq reads were mapped to exonic and intronic regions using annotations acquired from Ensembl GRCh38 version 87. DiffRAC was then used to decouple transcriptional and post-transcriptional effects and infer changes in mRNA stability.

### Animal Care

All animal protocols and procedures carried out were approved by the McGill University Animal Care Committee (UACC).

## Key Resources Table

| Antibody | Company | Catalog Number | Use |
| --- | --- | --- | --- |
| Anti-mouse ITGA7-647 | R&D systems | FAB3518R | FACS |
| Anti-mouse SCA1 (LY6A/E)-FITC | BD Pharmigen | 557405 | FACS |
| Anti-mouse CD31-PE | Invitrogen | 12-0311-81 | FACS |
| Anti-mouse CD45-BV785 | Biolegend | 103149 | FACS |
| Anti-mouse F4/80-APC-Cy7 | Biolegend | 123117 | FACS |
| Anti-mouse CD11b-PerCP-Cy5.5 | Biolegend | 101227 | FACS |
| PDGFRA | Cell Signaling | 31745 | IF |
| Anti-mouse F4/80 | Invitrogen | 12-4801-82 | IF |
| Anti-Mouse Pax7 | DSHB | AB_528428 | IF |

| Materials/Reagents | Company | Catalog Number | Use |
| --- | --- | --- | --- |
| Hoechst 33342 | Molecular Probes | H1399 | FACS |
| Goat Serum | Life Technologies | 500622 | IF |
| Prolong Gold Antifade with DAPI | Invitrogen | P3695 | IF |
| Collagenase D | Roche | 11088882001 | FACS |
| Dispase II | Roche | 04942078001 | FACS |
| Fetal Bovine Serum | Wisent | 080450 | FACS |
| PBS | Wisent | 311-425-CL | FACS |
| Bovine Serum Albumin | Sigma | A8022 | IF |
| Sucrose | Sigma | S7903 | IF |
| Isopentane | Sigma | M32631 | IF |
| UltraComp eBeads Compensation Beads | Invitrogen | 01-2222-41 | FACS |
| 40µm Nylon Cell Strainer | Falcon | C352340 | FACS |

| Commercial Kit | Company | Catalog Number |
| --- | --- | --- |
| SMART-Seq HT | Molecular Probes | 634456 |
| Nextera XT DNA Library Preparation Kit | Illumina | FC-131-1024/FC-131-1001 |
| NxSeq AmpFREE low DNA Library Kit | Lucigen | 14000 |
| EZ-96 DNA Methylation-Gold™ MagPrep kit | Zymo Research | D5042 |
| Quant-iT PicoGreen dsDNA Assay Kit | ThermoFisher | P7589 |

| Experimental Models: Mice | Company/Reference | Catalog Number |
| --- | --- | --- |
| C57BL/6J | Jackson Laboratory | 000664 |
| Tg(Pax7-EGFP)#Tajb (Pax7-nGFP) | (Sambasivan et al., 2009) | N/A |
| NOD-Prkdcem26Cd52Il2rgem26Cd22/NjuCrl (NCG) | Charles River | 572 |

## Author contributions

Conceptualization: VDS; Methodology: FL, VDS, RF, KS, AHC, HSN; Investigation: VDS, FL, RF, KS, DB, AHC, HSN, GP, TP; Visualization: FL, VDS, RF, HSN, AHC; Funding acquisition: VDS, HSN; Project administration: VDS; Supervision: VDS, HSN; Resources: VDS, AJA, JR, CC, HSN; Software: RF, AHC, TP, GP, HSN; Writing – original draft: FL, VDS; Writing – review & editing: FL, VDS, RF, DB, KS, AJA, HSN

## Acknowledgments

We thank Christian Young at the Lady Davis Institute for Medical Research—Jewish General Hospital—core facility for help with fluorescence-activated cell sorting (FACS). We thank Dr. Shahragim Tajbakhsh from Institut Pasteur, Paris, France for kindly providing us with Pax7-nGFP mice. This work was supported by a Disease team grant from the Stem Cell Network, a Natural Sciences and Engineering Research Council (NSERC) Discovery Grant and Richard & Edith Strauss Canada Foundation to VDS, as well as NSERC Discovery Grant (RGPIN-2018-05962) and Compute Canada resource allocations to HSN. GP is supported by training scholarships from the Canadian Institutes of Health Research (CIHR), the Fonds de recherche du Québec – Santé (FRQS), and Oncopole. HSN holds a Canada Research Chair funded by the CIHR.

## Data and availability

The Next-Generation Sequencing (NGS) data reported in this manuscript are available through the NCBI GEO super series GSE171534.

## Supplemental Information

### Supplemental Figures

**Figure S1.**
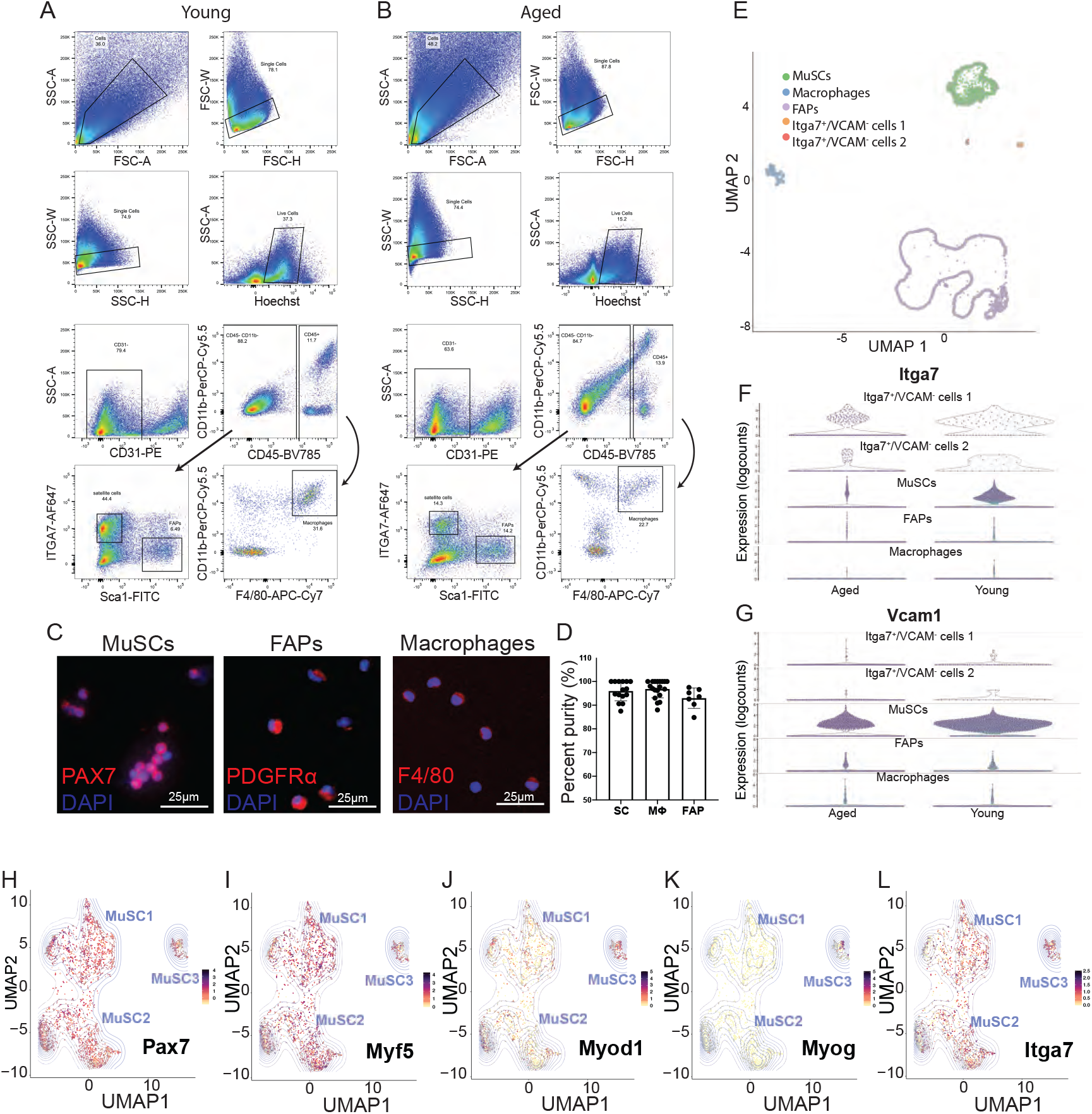
Fluorescence Activated Cell Sorting (FACS) Strategy for the Simultaneous Isolation of Pure Populations of MuSCs, Fibro-Adipogenic Progenitor (FAPs) Cells and Muscle-Resident Macrophages, Related to Figure 1. **(A-B)** Complete FACS sorting strategy for the simultaneous isolation of MuSCs (ITGA7^+^, CD31^-^, SCA1^-^, CD45^-^, CD11b-)(Pasut et al., 2012), macrophages (F4/80^+^, CD45^+^, CD11b^+^, CD31^-^)(Farup et al., 2015), and FAPs (SCA1^+^, ITGA7^-^, CD45^-^, CD11b^-^, CD31^-^)(Low et al., 2017) from young (A) and aged (B) mice. **(C)** Immunofluorescent staining of freshly-sorted MuSCs, macrophages and FAPs with selected representative markers (PAX7, PDGFRA, F4/80, respectively) of each population. (Scale bar = 25μm) **(D)** Quantification of the percentage of pure freshly-isolated cells based on immunofluorescence of PAX7 in MuSCs (n=1 isolation, n=15 fields of view, n=440 cells counted), PDGFRA in FAPs (n=2 isolations, n=7 fields of view, n=820 cells counted) and F4/80 in macrophages (n=2 isolations, n=19 fields of view, n=656 cells counted). **(E)** UMAP plot of young and aged MuSC, FAP and macrophage scRNA-seq samples before filtering out two ITGA7^+^/VCAM1^-^ cell populations. **(F-G)** Violin plots showing the expression of Itga7 (F) and Vcam1 (G) in young and aged MuSC, FAP and macrophage scRNA-seq samples before filtering out two ITGA7^+^/VCAM1^-^ cell populations. **(H-L)** Gene expression plots in the UMAP embedding for muscle-specific genes (*Pax7, Myf5, Myod1, Myog, Itga7*) in young and aged MuSCs.

**Figure S2.**
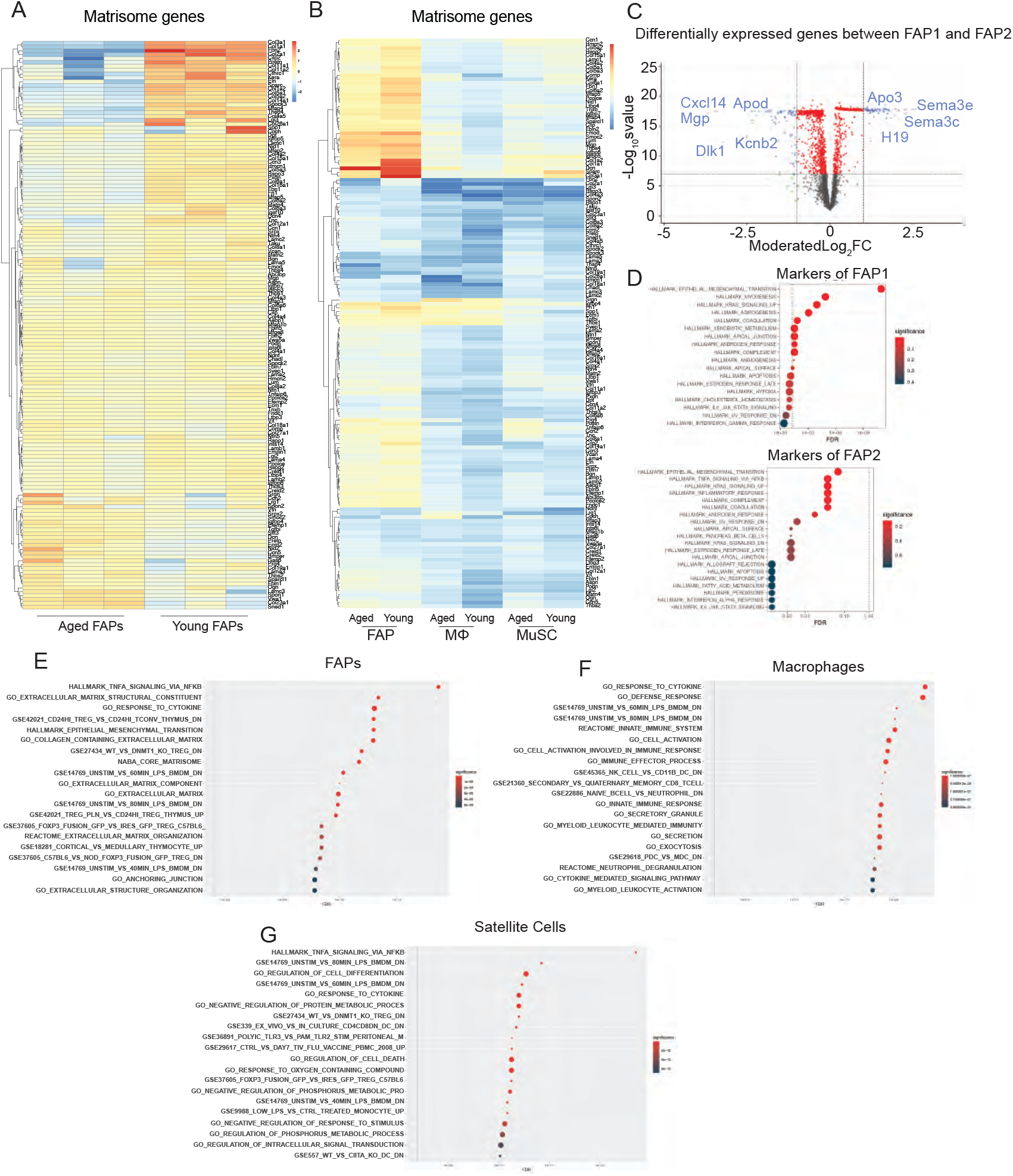
Genes Involved in Processes Such as Cytokine Signaling and Extracellular Matrix Composition are Perturbed in Aged MuSCs and Niche Cells, Related to Figure 1. **(A)** Heat map of matrisome genes in young and aged FAPs (color scale denotes beta-coefficients. **(B)** Heat map of matrisome genes (Naba et al., 2012) in young and aged macrophages, FAPs, and MuSCs. (color scale represents beta-coefficients) **(C)** Volcano plot of moderated LFC and s-value of the contract between FAP2 and FAP1 clusters. Positive values show genes upregulated in FAP2 subpopulation. **(D)** Gene set enrichment analysis of positive markers of FAP1 and FAP2 subpopulations using Hallmark gene sets (Liberzon et al., 2015). **(E)** Gene set enrichment analysis of young and aged FAPs using Hallmark gene sets. **(F)** Gene set enrichment analysis of young and aged macrophages using Hallmark gene sets. **(G)** Gene set enrichment analysis of young and aged MuSCs using Hallmark gene sets.

**Figure S3.**
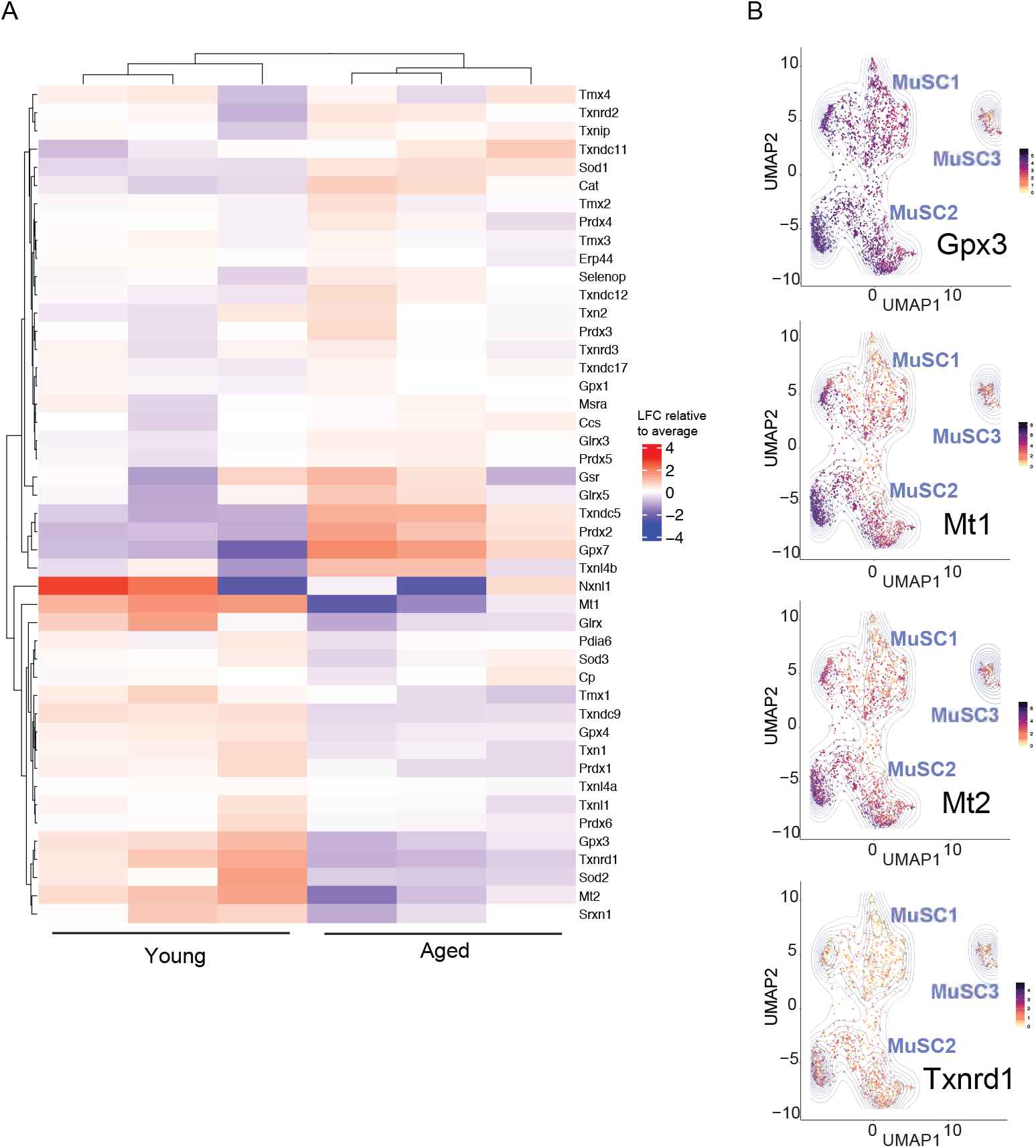
Aged MuSCs Display an Altered Expression Profile of Stress Response Genes, Related to Figure 2. **(A)** Heat map showing the LFC relative to the average expression of stress-response genes (Gelain et al., 2009) in young compared to aged MuSCs. **(B)** Gene expression plots in the UMAP embedding for select stress-response genes thar are differentially expressed between young and aged MuSCs.

**Figure S4.**
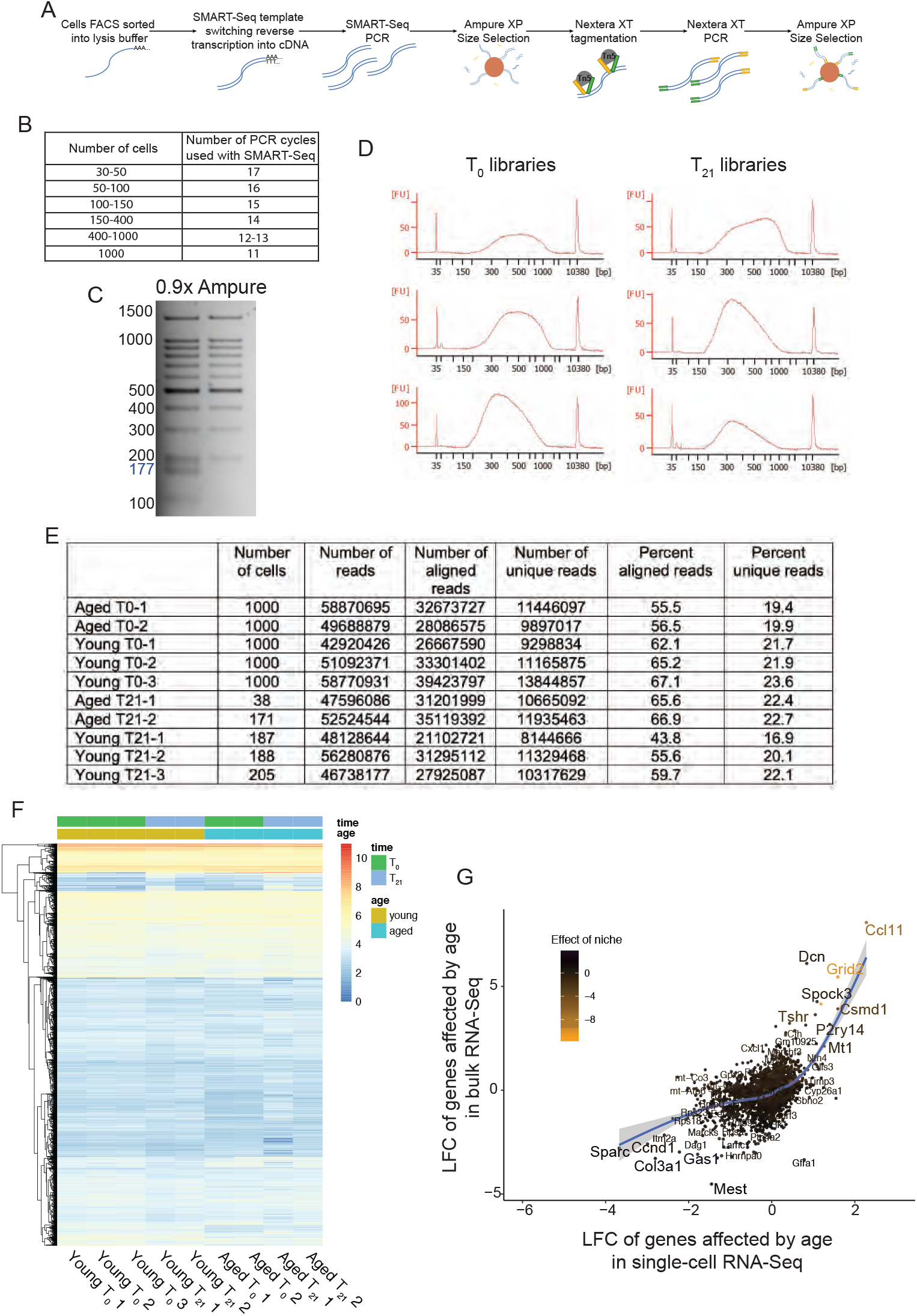
SMART-seq Technology Allows the Generation of RNA-sequencing Libraries from Rare Cell Types, Related to Figure 3. **(A)** Schematic diagram of library preparation method using SMART-seq (Picelli et al., 2014b) and Nextera XT(Adey et al., 2010). Briefly, MuSCs are sorted using FACS directly into SMART-seq reaction buffer, followed by template switching reverse transcription and PCR. After Ampure XP size selection at a 1:1 (v:v) ratio, cDNA is quantified and 0.15ng is used for Nextera XT tagmentation. After PCR addition of Illumina sequencing adapters, libraries are once again purified using Ampure XP at a 1:0.9 (v:v) ratio and sequenced. **(B)** Number of PCR cycles used with SMART-seq depending on the number of cells sorted by FACS. **(C)** Representative photo of Ampure XP size selection at a 0.9x ratio on a DNA ladder, visualized on an agarose gel, demonstrating its utility at removing primer dimers of <200bp. **(D)** Example of bioanalyzer profiles of SMART-seq libraries created from 50-1000 cells **(E)** Table of sequencing read information for SMART-seq libraries generated from different numbers of cells **(F)** Heat map of the top 6000 expressed genes in young and aged MuSC samples, at T_0_ and at T_21_, showing consistency within replicates. **(G)** Scatter plot of the LFCs of the effect of age from bulk RNA-seq and from scRNA-seq showing high concordance of differentially expressed genes from both modalities. Coloring is based on the size of the LFC effect of the allogeneic niche in bulk RNA-seq.

**Figure S5.**
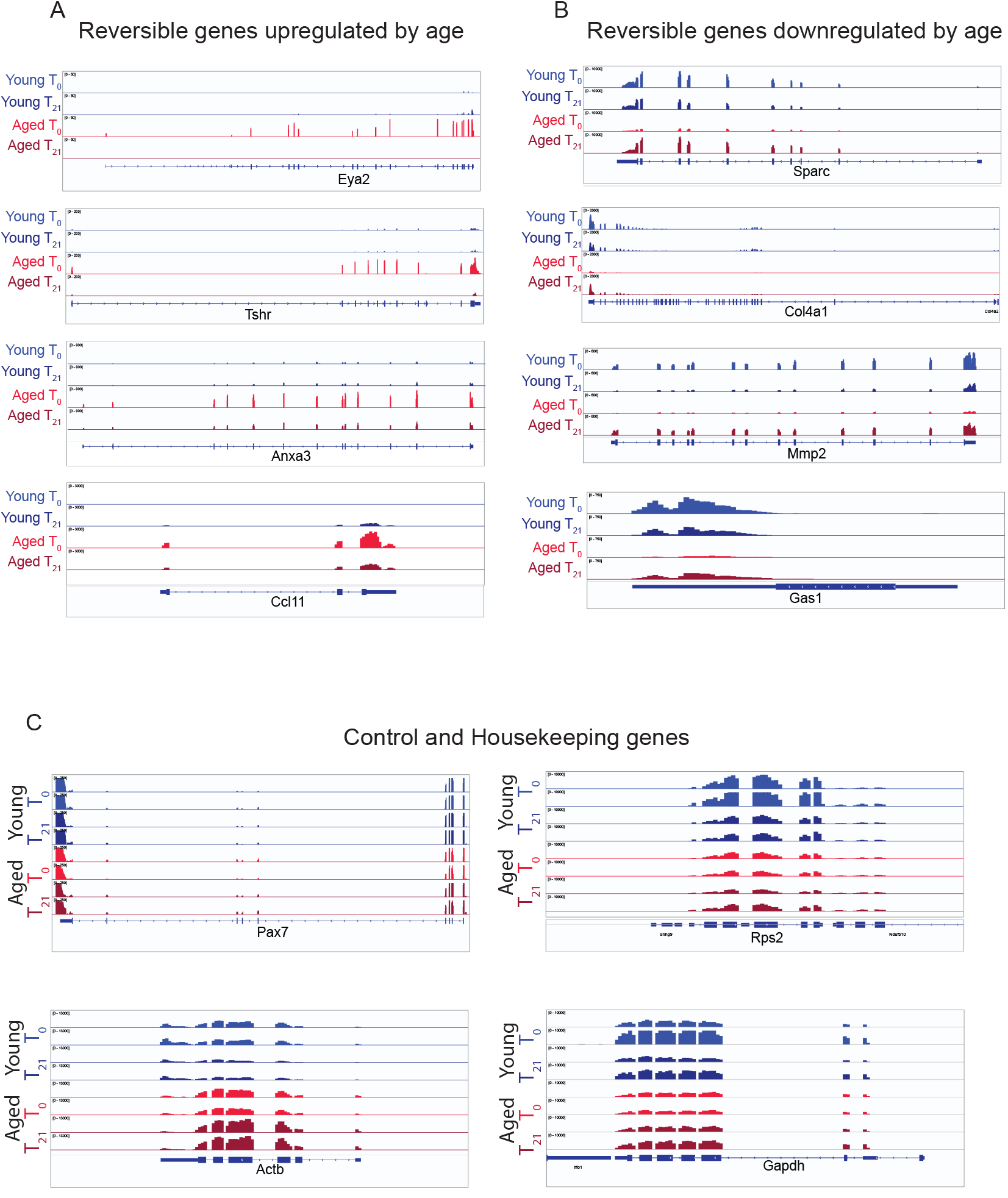
A Number of Genes in Aged MuSCs can be Restored to a Youthful Condition after Transplantation into a Young Niche, Related to Figure 4. **(A-B)** Representative IGV tracks of selected age-upregulated (A) or downregulated (B) genes (*Sparc, Col4a1, Mmp2, Eya2, Gas1, Prdx2, Tshr*) that are reversible to a youthful state by exposure to the young niche in aged MuSCs. **(C)** Representative IGV tracks of select control and housekeeping genes (*Pax7, Gapdh, Actb, Rps2*) in young and aged MuSCs from SMART-seq analysis at T_0_ and T_21_.

**Figure S6.**
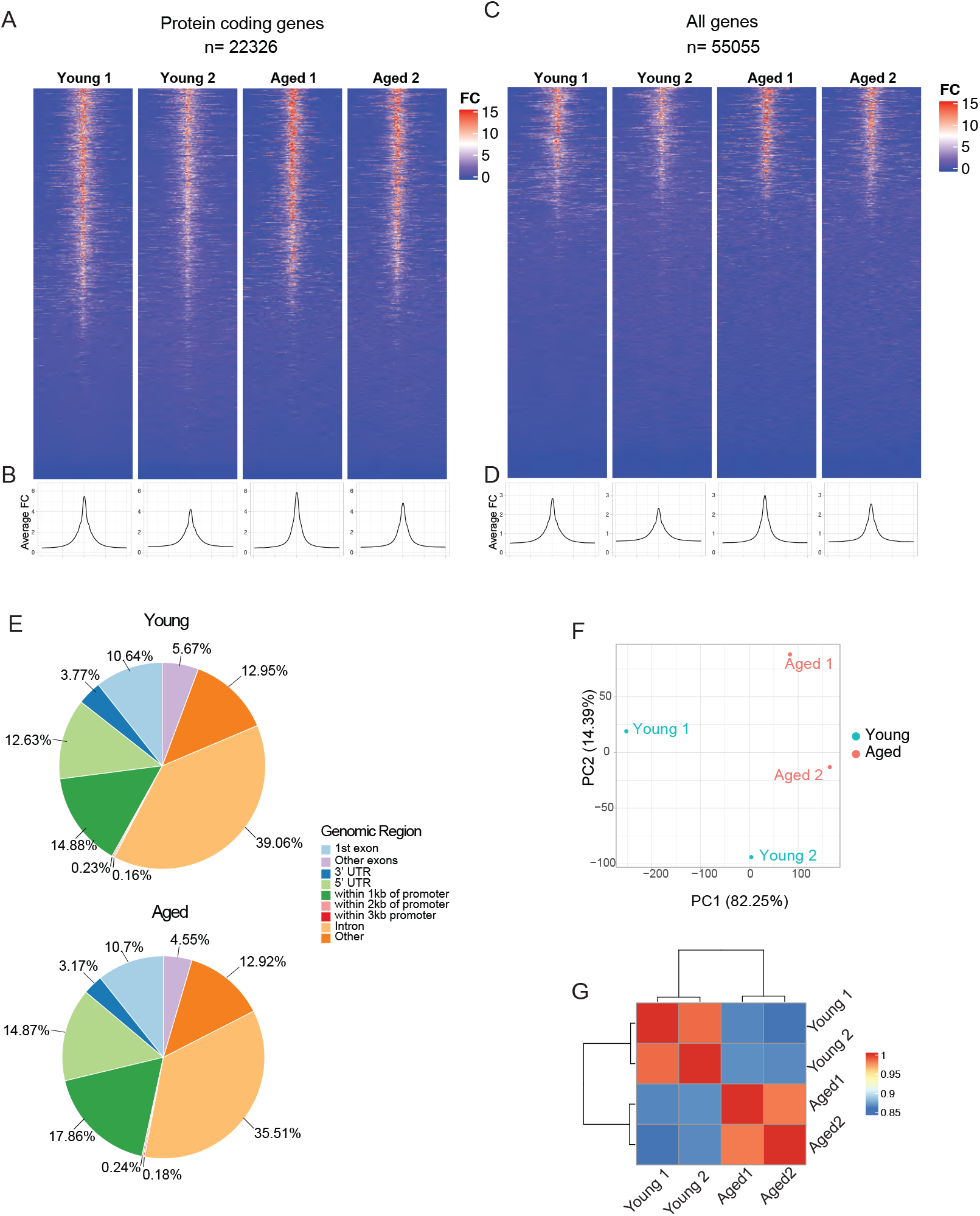
Analysis of Chromatin State by ATAC-seq from Young and Aged MuSCs, Related to Figure 5. **(A)** Pileup of ATAC-seq reads around the TSS of protein-coding genes in young and aged MuSCs. **(B)** Aggregation plots (Averageograms) showing the position of peaks around the TSS for protein-coding genes in young and aged MuSCs **(C)** Pileup of ATAC-seq reads around the TSS of all genes in young and aged MuSCs. **(D)** Aggregation plots (Averageograms) showing the position of peaks around the TSS for all genes in young and aged MuSCs **(E)** Pie charts showing proportions of ATAC-seq peaks throughout the genome in young and aged MuSC samples (n=2 biological replicates per group) **(F)** Principal Component Analysis (PCA) of young and aged MuSCs by ATAC-seq (n=2 biological replicates per group). **(G)** Heat map showing the level of similarity between young and aged MuSC ATAC-seq samples (n=2).

**Figure S7.**
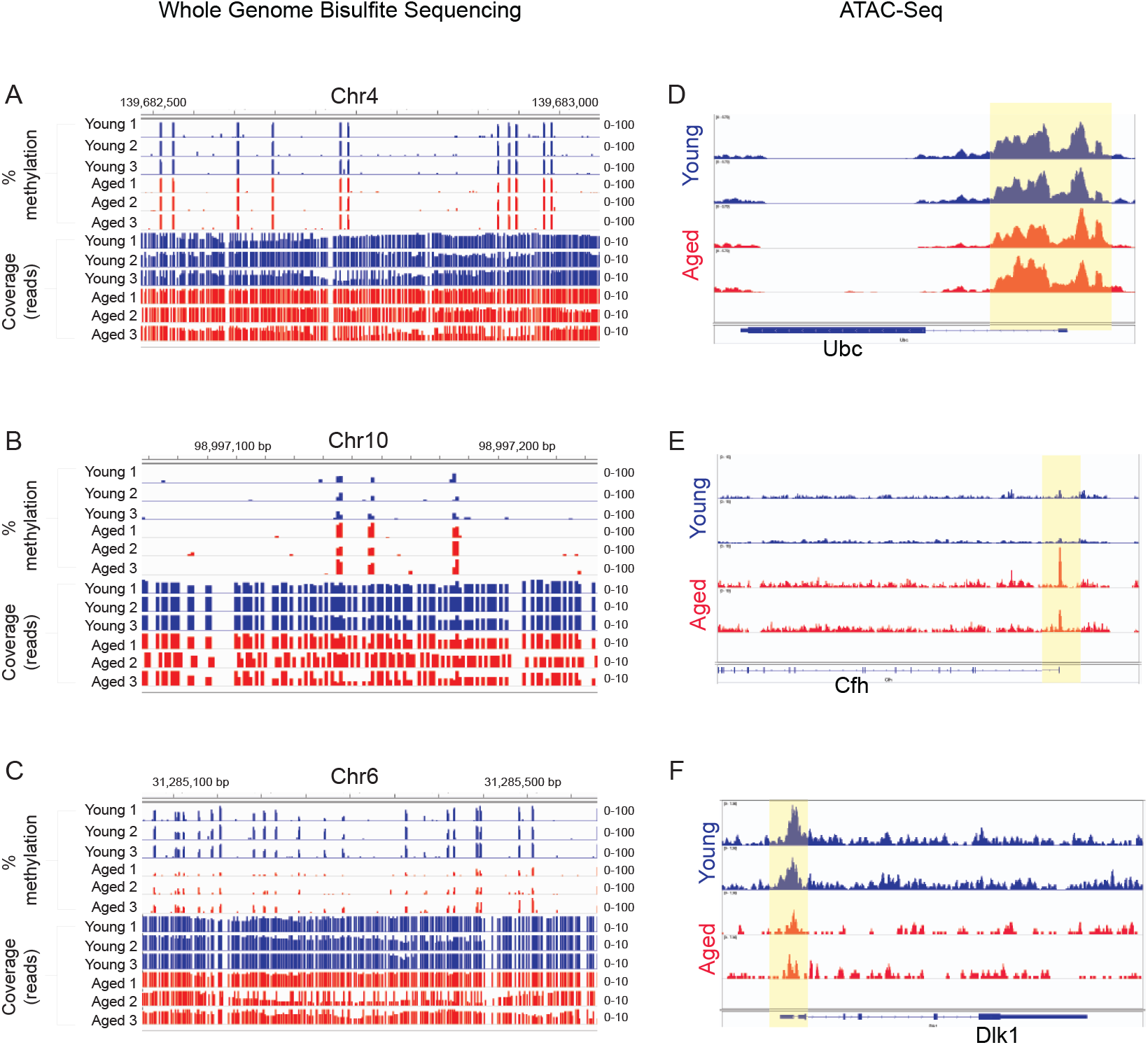
Chromatin is a Driver of the Age-Related Changes in MuSCs, Related to Figure 5. **(A-C)** Representative IGV tracks showing examples of chromosomal regions of equal DNA methylation (A), increased methylation (B), and decreased methylation (C) in aged compared to young MuSCs. **(D-F)** Representative IGV tracks of ATAC-seq reads for select example genes in young and aged MuSCs showing equal chromatin accessibility (D), increased accessibility (E), and decreased accessibility (F) In aging. Yellow boxes indicate peaks close to the TSS of each gene.

## References and Notes

Adey, A., Morrison, H.G., Asan Xun, X., Kitzman, J.O., Turner, E.H., Stackhouse, B., MacKenzie, A.P., Caruccio, N.C., Zhang, X., et al. (2010). Rapid, low-input, low-bias construction of shotgun fragment libraries by high-density in vitro transposition. Genome biology 11, R119.

Ancel, S., Mashinchian, O., and Feige, J.N. (2019). Adipogenic progenitors keep muscle stem cells young. Aging 11, 7331–7333.

Angeloni, A., and Bogdanovic, O. (2019). Enhancer DNA methylation: implications for gene regulation. Essays in biochemistry 63, 707–715.

Benjamini, Y., and Hochberg, Y. (1995). Controlling the False Discovery Rate: A Practical and Powerful Approach to Multiple Testing. Journal of the Royal Statistical Society: Series B (Methodological) 57, 289–300.

Berger, M.J., and Doherty, T.J. (2010). Sarcopenia: prevalence, mechanisms, and functional consequences. Interdisciplinary topics in gerontology 37, 94–114.

Bernet, J.D., Doles, J.D., Hall, J.K., Kelly Tanaka, K., Carter, T.A., and Olwin, B.B. (2014). p38 MAPK signaling underlies a cell-autonomous loss of stem cell self-renewal in skeletal muscle of aged mice. Nat Med 20, 265–271.

Biferali, B., Proietti, D., Mozzetta, C., and Madaro, L. (2019). Fibro–Adipogenic Progenitors Cross-Talk in Skeletal Muscle: The Social Network. Frontiers in Physiology 10.

Blackburn, D.M., Lazure, F., Corchado, A.H., Perkins, T.J., Najafabadi, H.S., and Soleimani, V.D. (2019). High-Resolution Genome-Wide Expression Analysis of Single Myofibers Using SMART-Seq. Journal of Biological Chemistry.

Blattler, A., Yao, L., Witt, H., Guo, Y., Nicolet, C.M., Berman, B.P., and Farnham, P.J. (2014). Global loss of DNA methylation uncovers intronic enhancers in genes showing expression changes. Genome biology 15, 469.

Catoni, M., Tsang, J.M., Greco, A.P., and Zabet, N.R. (2018). DMRcaller: a versatile R/Bioconductor package for detection and visualization of differentially methylated regions in CpG and non-CpG contexts. Nucleic Acids Res 46, e114.

Chakkalakal, J.V., Jones, K.M., Basson, M.A., and Brack, A.S. (2012). The aged niche disrupts muscle stem cell quiescence. Nature 490, 355–360.

Conboy, I.M., Conboy, M.J., Wagers, A.J., Girma, E.R., Weissman, I.L., and Rando, T.A. (2005). Rejuvenation of aged progenitor cells by exposure to a young systemic environment. Nature 433, 760–764.

Corces, M.R., Trevino, A.E., Hamilton, E.G., Greenside, P.G., Sinnott-Armstrong, N.A., Vesuna, S., Satpathy, A.T., Rubin, A.J., Montine, K.S., Wu, B., et al. (2017). An improved ATAC-seq protocol reduces background and enables interrogation of frozen tissues. Nature Methods 14, 959–962.

Cruz-Jentoft, A.J., Bahat, G., Bauer, J., Boirie, Y., Bruyère, O., Cederholm, T., Cooper, C., Landi, F., Rolland, Y., Sayer, A.A., et al. (2019). Sarcopenia: revised European consensus on definition and diagnosis. Age Ageing 48, 16–31.

Cui, C.-Y., Driscoll, R.K., Piao, Y., Chia, C.W., Gorospe, M., and Ferrucci, L. (2019). Skewed macrophage polarization in aging skeletal muscle. Aging cell 18, e13032–e13032.

Di Raimo, T., Leopizzi, M., Mangino, G., Rocca, C.D., Businaro, R., Longo, L., and Lo Vasco, V.R. (2016). Different expression and subcellular localization of Phosphoinositide-specific Phospholipase C enzymes in differently polarized macrophages. J Cell Commun Signal 10, 283–293.

El Haddad, M., Jean, E., Turki, A., Hugon, G., Vernus, B., Bonnieu, A., Passerieux, E., Hamade, A., Mercier, J., Laoudj-Chenivesse, D., et al. (2012). Glutathione peroxidase 3, a new retinoid target gene, is crucial for human skeletal muscle precursor cell survival. J Cell Sci 125, 6147–6156.

Farup, J., Madaro, L., Puri, P.L., and Mikkelsen, U.R. (2015). Interactions between muscle stem cells, mesenchymal-derived cells and immune cells in muscle homeostasis, regeneration and disease. Cell Death & Disease 6, e1830–e1830.

Federico, A., and Monti, S. (2020). hypeR: an R package for geneset enrichment workflows. Bioinformatics 36, 1307–1308.

Garg, K., and Boppart, M.D. (2016). Influence of exercise and aging on extracellular matrix composition in the skeletal muscle stem cell niche. J Appl Physiol (1985) 121, 1053–1058.

Gelain, D.P., Dalmolin, R.J., Belau, V.L., Moreira, J.C., Klamt, F., and Castro, M.A. (2009). A systematic review of human antioxidant genes. Frontiers in bioscience (Landmark edition) 14, 4457–4463.

Gilbert, P.M., Havenstrite, K.L., Magnusson, K.E.G., Sacco, A., Leonardi, N.A., Kraft, P., Nguyen, N.K., Thrun, S., Lutolf, M.P., and Blau, H.M. (2010). Substrate elasticity regulates skeletal muscle stem cell self-renewal in culture. Science 329, 1078–1081.

Haghverdi, L., Lun, A.T.L., Morgan, M.D., and Marioni, J.C. (2018). Batch effects in single-cell RNA-sequencing data are corrected by matching mutual nearest neighbors. Nat Biotechnol 36, 421–427.

Hernando-Herraez, I., Evano, B., Stubbs, T., Commere, P.-H., Jan Bonder, M., Clark, S., Andrews, S., Tajbakhsh, S., and Reik, W. (2019). Ageing affects DNA methylation drift and transcriptional cell-to-cell variability in mouse muscle stem cells. Nature Communications 10, 4361.

Joe, A.W., Yi, L., Natarajan, A., Le Grand, F., So, L., Wang, J., Rudnicki, M.A., and Rossi, F.M. (2010). Muscle injury activates resident fibro/adipogenic progenitors that facilitate myogenesis. Nat Cell Biol 12, 153–163.

Kimmel, J.C., Hwang, A.B., Scaramozza, A., Marshall, W.F., and Brack, A.S. (2020). Aging induces aberrant state transition kinetics in murine muscle stem cells. Development, dev.183855.

Krueger, F., and Andrews, S.R. (2011). Bismark: a flexible aligner and methylation caller for Bisulfite-Seq applications. Bioinformatics 27, 1571–1572.

Langmead, B., and Salzberg, S.L. (2012). Fast gapped-read alignment with Bowtie 2. Nature Methods 9, 357–359.

Lee, B.C., Lee, S.-G., Choo, M.-K., Kim, J.H., Lee, H.M., Kim, S., Fomenko, D.E., Kim, H.-Y., Park, J.M., and Gladyshev, V.N. (2017). Selenoprotein MsrB1 promotes anti-inflammatory cytokine gene expression in macrophages and controls immune response in vivo. Scientific Reports 7, 5119.

Li, H., Chen, Q., Li, C., Zhong, R., Zhao, Y., Zhang, Q., Tong, W., Zhu, D., and Zhang, Y. (2019). Muscle-secreted granulocyte colony-stimulating factor functions as metabolic niche factor ameliorating loss of muscle stem cells in aged mice. The EMBO Journal 38, e102154.

Liberzon, A., Birger, C., Thorvaldsdóttir, H., Ghandi, M., Mesirov, J.P., and Tamayo, P. (2015). The Molecular Signatures Database (MSigDB) hallmark gene set collection. Cell Syst 1, 417–425.

Lombard, D.B., Chua, K.F., Mostoslavsky, R., Franco, S., Gostissa, M., and Alt, F.W. (2005). DNA Repair, Genome Stability, and Aging. Cell 120, 497–512.

Love, M.I., Huber, W., and Anders, S. (2014). Moderated estimation of fold change and dispersion for RNA-seq data with DESeq2. Genome biology 15, 550.

Low, M., Eisner, C., and Rossi, F. (2017). Fibro/Adipogenic Progenitors (FAPs): Isolation by FACS and Culture. In Muscle Stem Cells: Methods and Protocols, E. Perdiguero, and D.D.W. Cornelison, eds. (New York, NY: Springer New York), pp. 179–189.

Lukjanenko, L., Jung, M.J., Hegde, N., Perruisseau-Carrier, C., Migliavacca, E., Rozo, M., Karaz, S., Jacot, G., Schmidt, M., Li, L., et al. (2016). Loss of fibronectin from the aged stem cell niche affects the regenerative capacity of skeletal muscle in mice. Nat Med 22, 897–905.

Lukjanenko, L., Karaz, S., Stuelsatz, P., Gurriaran-Rodriguez, U., Michaud, J., Dammone, G., Sizzano, F., Mashinchian, O., Ancel, S., Migliavacca, E., et al. (2019). Aging Disrupts Muscle Stem Cell Function by Impairing Matricellular WISP1 Secretion from Fibro-Adipogenic Progenitors. Cell Stem Cell 24, 433-446.e437.

Martin, M. (2011). Cutadapt removes adapter sequences from high-throughput sequencing reads. 2011 17, 3.

McInnes, L., Healy, J., and Melville, J. (2018). UMAP: Uniform Manifold Approximation and Projection for Dimension Reduction, pp. 1802.03426.

Melouane, A., Carbonell, A., Yoshioka, M., Puymirat, J., and St-Amand, J. (2018). Implication of SPARC in the modulation of the extracellular matrix and mitochondrial function in muscle cells. PLOS ONE 13, e0192714.

Morgan, J.E., Hoffman, E.P., and Partridge, T.A. (1990). Normal myogenic cells from newborn mice restore normal histology to degenerating muscles of the mdx mouse. J Cell Biol 111, 2437–2449.

Morgan, J.E., Pagel, C.N., Sherratt, T., and Partridge, T.A. (1993). Long-term persistence and migration of myogenic cells injected into pre-irradiated muscles of mdx mice. Journal of the neurological sciences 115, 191–200.

Mourikis, P., Sambasivan, R., Castel, D., Rocheteau, P., Bizzarro, V., and Tajbakhsh, S. (2012). A critical requirement for notch signaling in maintenance of the quiescent skeletal muscle stem cell state. Stem cells (Dayton, Ohio) 30, 243–252.

Naba, A., Clauser, K.R., Hoersch, S., Liu, H., Carr, S.A., and Hynes, R.O. (2012). The matrisome: in silico definition and in vivo characterization by proteomics of normal and tumor extracellular matrices. Molecular & cellular proteomics : MCP 11, M111.014647.

Ocampo, A., Reddy, P., Martinez-Redondo, P., Platero-Luengo, A., Hatanaka, F., Hishida, T., Li, M., Lam, D., Kurita, M., Beyret, E., et al. (2016). In Vivo Amelioration of Age-Associated Hallmarks by Partial Reprogramming. Cell 167, 1719-1733.e1712.

Oh, J., Lee, Y.D., and Wagers, A.J. (2014). Stem cell aging: mechanisms, regulators and therapeutic opportunities. Nat Med 20, 870–880.

Oh, J., Sinha, I., Tan, K.Y., Rosner, B., Dreyfuss, J.M., Gjata, O., Tran, P., Shoelson, S.E., and Wagers, A.J. (2016). Age-associated NF-κB signaling in myofibers alters the satellite cell niche and re-strains muscle stem cell function. Aging (Albany NY) 8, 2871–2896.

Parisi, L., Gini, E., Baci, D., Tremolati, M., Fanuli, M., Bassani, B., Farronato, G., Bruno, A., and Mortara, L. (2018). Macrophage Polarization in Chronic Inflammatory Diseases: Killers or Builders? Journal of Immunology Research 2018, 8917804.

Pasut, A., Oleynik, P., and Rudnicki, M.A. (2012). Isolation of muscle stem cells by fluorescence activated cell sorting cytometry. Methods in molecular biology (Clifton, NJ) 798, 53–64.

Patro, R., Duggal, G., Love, M.I., Irizarry, R.A., and Kingsford, C. (2017). Salmon provides fast and bias-aware quantification of transcript expression. Nature Methods 14, 417–419.

Perron, G., Jandaghi, P., Rajaee, M., Alkallas, R., Riazalhosseini, Y., and Najafabadi, H.S. (2021). Pan-cancer analysis of mRNA stability for decoding tumour post-transcriptional programs. bioRxiv, 2020.2012.2030.424872.

Picelli, S., Björklund, A.K., Reinius, B., Sagasser, S., Winberg, G., and Sandberg, R. (2014a). Tn5 transposase and tagmentation procedures for massively scaled sequencing projects. Genome Res 24, 2033–2040.

Picelli, S., Faridani, O.R., Bjorklund, A.K., Winberg, G., Sagasser, S., and Sandberg, R. (2014b). Full-length RNA-seq from single cells using Smart-seq2. Nat Protoc 9, 171–181.

Pons, P., and Latapy, M. (2005). Computing Communities in Large Networks Using Random Walks. Paper presented at: Computer and Information Sciences-ISCIS 2005 (Berlin, Heidelberg: Springer Berlin Heidelberg).

Quinlan, A.R. (2014). BEDTools: The Swiss-Army Tool for Genome Feature Analysis. Current protocols in bioinformatics 47, 11.12.11-34.

Qunhua, L., James, B.B., Haiyan, H., and Peter, J.B. (2011). Measuring reproducibility of high-throughputexperiments. The Annals of Applied Statistics 5, 1752–1779.

Risso, D., Perraudeau, F., Gribkova, S., Dudoit, S., and Vert, J.-P. (2018). A general and flexible method for signal extraction from single-cell RNA-seq data. Nature Communications 9, 284.

Sambasivan, R., Gayraud-Morel, B., Dumas, G., Cimper, C., Paisant, S., Kelly, R.G., and Tajbakhsh, S. (2009). Distinct Regulatory Cascades Govern Extraocular and Pharyngeal Arch Muscle Progenitor Cell Fates. Developmental Cell 16, 810–821.

Sampath, S.C., Sampath, S.C., Ho, A.T.V., Corbel, S.Y., Millstone, J.D., Lamb, J., Walker, J., Kinzel, B., Schmedt, C., and Blau, H.M. (2018). Induction of muscle stem cell quiescence by the secreted niche factor Oncostatin M. Nature Communications 9, 1531.

Shefer, G., Van de Mark, D.P., Richardson, J.B., and Yablonka-Reuveni, Z. (2006). Satellite-cell pool size does matter: Defining the myogenic potency of aging skeletal muscle. Developmental Biology 294, 50–66.

Soneson, C., Love, M.I., and Robinson, M.D. (2015). Differential analyses for RNA-seq: transcript-level estimates improve gene-level inferences. F1000Research 4, 1521.

Soneson, C., Srivastava, A., Patro, R., and Stadler, M.B. (2021). Preprocessing choices affect RNA velocity results for droplet scRNA-seq data. PLOS Computational Biology 17, e1008585.

Srivastava, A., Malik, L., Sarkar, H., Zakeri, M., Almodaresi, F., Soneson, C., Love, M.I., Kingsford, C., and Patro, R. (2020). Alignment and mapping methodology influence transcript abundance estimation. Genome biology 21, 239.

Srivastava, A., Malik, L., Smith, T., Sudbery, I., and Patro, R. (2019). Alevin efficiently estimates accurate gene abundances from dscRNA-seq data. Genome biology 20, 65.

Stephens, M. (2017). False discovery rates: a new deal. Biostatistics (Oxford, England) 18, 275–294.

Takahashi, K., and Yamanaka, S. (2006). Induction of Pluripotent Stem Cells from Mouse Embryonic and Adult Fibroblast Cultures by Defined Factors. Cell 126, 663–676.

Villeda, S.A., Luo, J., Mosher, K.I., Zou, B., Britschgi, M., Bieri, G., Stan, T.M., Fainberg, N., Ding, Z., Eggel, A., et al. (2011). The ageing systemic milieu negatively regulates neurogenesis and cognitive function. Nature 477, 90–94.

Wang, Y., Wehling-Henricks, M., Samengo, G., and Tidball, J.G. (2015). Increases of M2a macrophages and fibrosis in aging muscle are influenced by bone marrow aging and negatively regulated by muscle-derived nitric oxide. Aging Cell 14, 678–688.

Zhu, A., Ibrahim, J.G., and Love, M.I. (2019). Heavy-tailed prior distributions for sequence count data: removing the noise and preserving large differences. Bioinformatics 35, 2084–2092.

Zismanov, V., Chichkov, V., Colangelo, V., Jamet, S., Wang, S., Syme, A., Koromilas Antonis E., and Crist, C. (2016). Phosphorylation of eIF2α Is a Translational Control Mechanism Regulating Muscle Stem Cell Quiescence and Self-Renewal. Cell Stem Cell 18, 79–90.

